# Feedback regulation between FOXM1 and APC/C^Cdh1^ determines the changes in cell cycle dynamics during aging

**DOI:** 10.1101/2025.01.26.634919

**Authors:** Ana Rita Araújo, Filipa Gaspar-Silva, Elsa Logarinho

## Abstract

Aging is characterised by a loss of regenerative capacity, though it remains elusive how aged proliferating cells slowdown cycling eventually becoming senescent. We previously found that repression of the FOXM1 transcription factor accounts for mitotic decline during aging due to a global transcriptional shutdown of mitotic genes in proliferating cells. Intriguingly, a 1.5-fold increase in both cell cycle and mitosis durations was observed in elderly cells in deviancy to a previous study showing mitosis to be temporally insulated from variability in earlier cell cycle phases due to the robustness of the positive feedback loop controlling CDK1-Cyclin B1 activity. Thus, we asked if molecular thresholds controlling cell cycle phase transitions become unfitted with aging. Here, we used live-cell imaging of primary human dermal fibroblasts of advancing age donors in combination with high-throughput image analysis, to investigate age-related changes in cell cycle dynamics. Interestingly, we found mitosis insularity to be gradually lost along aging due to defective switch-like activation of CDK1 at mitotic entry driven by *FOXM1* repression. Moreover, we found the levels of FZR1/Cdh1 co-activator of APC/C, the E3-ubiquitin ligase directing the proteolytic degradation of FOXM1 at mitotic exit, to increase with advancing age. Importantly, *FZR1/Cdh1* repression was shown to restore cell cycle fitness and FOXM1 levels in aged proliferating cells, preventing the accumulation of cell cycle inhibitors and senescence markers in their progeny. Thus, changes in FOXM1 and APC/C^Cdh1^ interlinked activities account for the loss of proliferative capacity and senescence accrual during aging, thereby delivering useful markers and/or targets to explore in anti-aging approaches.

## INTRODUCTION

The unprecedented rate at which the world’s elderly population is growing, is leading to significant burden on the healthcare systems due to higher incidence of old-age co-morbidities^1–3^. Therefore, it is a priority to understand the aging process at the molecular, cellular, and physiological levels to come out with comprehensive strategies for healthspan extension.

The hallmarks of aging have been proposed and among them, cellular senescence, a state of permanent cell cycle arrest with several stereotypical features including the secretion of pro-inflammatory molecules (or SASP), has gained solid evidence for its contribution to aging and age-related diseases ^4,5^. However, the mechanisms behind cell cycle slowdown, error propensity, and ultimately, senescence onset in aged cells, remain poorly understood.

The decision to divide is a fundamental cellular response, and the networks that control cell division can adapt and remodel in different biological contexts. The cell cycle is characterized by a sequence of events – G1-, S-, G2-phases and Mitosis – by which a cell gives rise to two daughter cells ^6,7^. The fidelity of cell division depends on a complex regulatory network, which controls the timing and order of all cell cycle events. Even though the molecular machinery that drives cell division cycles is common and evolutionarily conserved in all tissues, it is known that distinct cell types exhibit very different cycles. In essence, progression through cell cycle is dependent on the activation and inactivation of cyclin-dependent kinase complexes (CDKs) and the switching between DNA duplication and DNA segregation in all cells ^6,7^. Moreover, biochemical feedback loops keep transitions through cell cycle ordered and unidirectional, adding an extra layer of control to the cell cycle network ^8,9^.

All events in mitosis are driven by the activation of the CDK1-Cyclin B1 complex that is heavily regulated by positive feedback control^10–12^. It activates its own activator – CDC25^13,14^– and inhibits its own inhibitor – WEE1^15–17^–, which creates a switch-like activation of CDK1-Cyclin B1 that turns mitotic entry sudden and irreversible. It was previously demonstrated that amongst different cell types (from embryonic stem cells to differentiated and cancer cells), mitotic duration is short, constant, and independent of variability in upstream cell cycle events. The mechanism underlying temporal insulation of mitosis is the positive feedback loop at mitotic entry, that when impaired, turns mitosis longer, variable and coupled with previous cell cycle events^18^. Also, positive feedback disruption in the mother cells affects the daughter cells subsequent cycles.

Interestingly, it was previously shown that human dermal fibroblasts (HDFs) retrieved from skin biopsies of elderly donors, exhibit cell cycle slowdown, increased mitotic duration and a higher rate of mitotic defects^19^. Repression of Forkhead Box M1 (FOXM1) transcriptional activity was disclosed as the mechanism behind mitotic decline during aging leading to a global shutdown of mitotic genes in older cells^19^. First CDK1-Cyclin A and later CDK1-Cyclin B1 complexes phosphorylate FOXM1 that is subsequently recruited by the MYB-MuvB (MMB) complex to trigger the expression of late cell cycle genes that will drive G2-phase and Mitosis ^20^. Moreover, FOXM1 activity is also under positive feedback regulation at G2/M transition, as FOXM1 is activated by CDK1-Cyclin B1 while it induces Cyclin B1 transcription to activate more CDK1-Cyclin B1 complexes^21,22^. By the end of mitosis, degradation of Cyclin B1 turns off CDK1-Cyclin B1 activity, allowing for accumulation of the E3 ubiquitin ligase, APC/C^Cdh^^1^, which drives FOXM1 proteasomal degradation ^23,24^.

FOXM1 repression has been causally linked to cellular senescence and aging. FOXM1 levels are decreased in fibroblasts from middle-age and elderly healthy donors, as well as from patients with Hutchinson-Gilford progeria, a rare genetic disorder characterized by premature aging symptoms at a young age^19,25^. Notably, FOXM1 overexpression was able to regress the expression of a comprehensive gene network driving senescence and SASP^19,25,26^. APC/C^Cdh1^ activity levels are kept low in embryonic stem cells with short division cycles and, in contrast, APC/C^Cdh1^ levels are high in differentiated cells as neurons that never re-enter the cell cycle^27,28^.

Here, we used live-cell imaging analysis of HDFs collected from healthy Caucasian males with ages ranging from neonatal to octogenarian ^19,25,29^ to ascertain for changes in cell cycle dynamics along natural aging and how this impacts proliferative fitness and senescence accrual. We found the cell cycle length to increase steadily along aging with all cell cycle phases showing extended duration. Temporal insulation of mitosis from previous cell cycle phases was found to be lost with aging due to a defective activation of CDK1-Cyclin B1 complex at mitotic entry induced by FOXM1 repression. Notably, both cell cycle dynamics and mitotic insularity were rescued in old cells by overexpressing FOXM1. Modulation of FZR/Cdh1 levels also induced changes in cell cycle dynamics, with Cdh1 overexpression leading to FOXM1 repression, cell cycle slowdown and senescence accrual in young cells, and with Cdh1 repression rescuing these features in aged cells. Importantly, FOXM1 and APC/C^Cdh1^ activities were shown to influence cell cycle fitness in the progeny, with prolonged mitosis in aged mother cells with low FOXM1/high Cdh1 imprinting their daughter cells for *CDKN2A/p16* expression and senescence propensity. Altogether, we disclose that changes in the feedback regulatory loop comprising CDK1, FOXM1 and APC/C^Cdh1^ account for age-related cell cycle frailty and senescence accrual.

## RESULTS

### Extended cell cycle phases and loss of mitotic temporal insularity with aging

To ascertain for changes in cell cycle dynamics along aging, HDFs retrieved from healthy donors with ages ranging from neonatal to octogenarian, and expressing the PIP-FUCCI sensor, were imaged for two consecutive divisions (Fig. 1A,B) ^30^. The duration of G1-, S-, G2- and M-phase was measured based on the appearance/disappearance of sensors and nuclear envelope breakdown (NEB) or reformation (NER) as shown in Fig. 1B. Total cell cycle length was measured as the time between two consecutive NER events. Interestingly, we found cell cycle duration to steadily increase with advancing age, showing 1.2-fold increase between every two consecutive age ranges (13.3 ± 2.2 hrs in neonatal, 16.5 ± 5.0 hrs in 10- year-old, 19.2 ± 7.5 hrs in 21-year-old, 22.3 ± 9.5 hrs in 42-year-old and 24.1 ± 10.0 hrs in 87-year-old) (Fig. 1C and Fig. S1A,B). We then asked if all cell cycle phases get delayed or if there is a specific phase that mainly accounts for cell cycle extension with aging. We found the duration of all cell cycle phases to increase with aging, including mitosis as previously reported (1.1-fold between each two consecutive ages)^19^, with G1-phase increasing the most (1.5-fold between each two consecutive ages) (Fig. 1D and Fig. S1A,B). Changes in cell cycle dynamics were highlighted by the correlation between the duration of the different cell cycle phases. Mitosis and G1-phase as well as S-phase and G2-phase start as inversely correlated (Pearson correlation coefficient, *r*) in young cells (neonatal HDFs: *r*=-0.14 and *r*=- 0.23, respectively) and become gradually correlated as the cells age (87-year-old HDFs: *r*=0.38 and *r*=0.32, respectively) (Fig. S1C). Correlations were not observed between G1-phase and S-phase (neonatal *r=*0.031 and 87-year-old *r*=0.11) (Fig. S1C).

**Figure 1.**
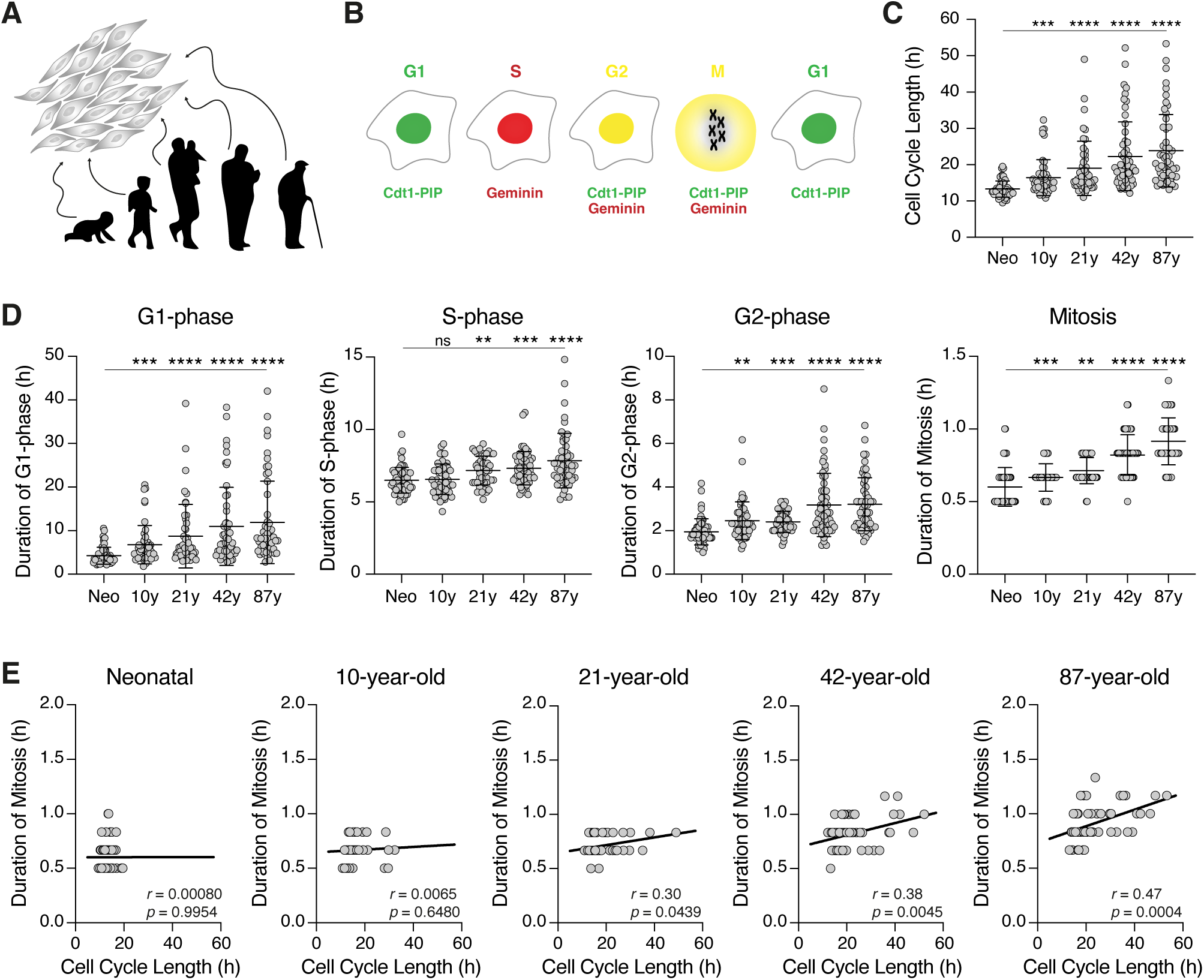
Mitotic duration becomes steadily correlated with cell cycle length along aging. ***A.*** *In vitro* model of cellular aging. Early passage human dermal fibroblast (HDF) cultures established from skin biopsies of different age donors. ***B.*** PIP-FUCCI genetically encoded fluorescence sensor that enables real-time monitoring of distinct cell cycle phases ^30^. ***C.*** Cell cycle length measured by the time between two consecutive mitoses in neonatal, 10-, 21-, 42- and 87-year-old HDFs expressing the PIP-FUCCI. Mean ± SD are shown as bars. *** *p* ≤0.001; **** *p* ≤0.0001 by Mann-Whitney U test, relative to neonatal HDFs. ***D.*** Duration of G1-phase, S-phase, G2-phase, and mitosis measured by single live-cell imaging in neonatal to 87-year-old HDFs expressing the PIP-FUCCI biosensor. Mean ± SD are shown as bars. ns nonsignificant; ** *p*≤0.01; *** *p*≤0.001; **** *p*≤0.0001 by Mann-Whitney U test, relative to neonatal HDFs. ***E.*** Duration of mitosis measured by the time between nuclear envelope breakdown (NEB) and nuclear envelope reformation (NER) as a function of cell cycle duration in neonatal to 87-year-old HDFs expressing the PIP-FUCCI. The trend lines, Pearson coefficient (*r*) and *p*-value are shown. n>50 cells, ≥2 independent experiments.

Since mitotic duration was found to increase with aging (Fig. 1D and Fig. S1A,B), we asked if the previously reported temporal insulation of mitosis^18^ is lost in older cells. Correlation analysis between cell cycle and mitotic durations revealed that in young HDFs mitosis was uncoupled from previous cell cycle events (*r=*0.001 in neonatal and *r=*0.07 in 10-year-old) (Fig. 1E). 21-year-old HDFs showed a mild correlation between cell cycle length and mitosis (*r*=0.30), and HDFs from older donors showed a striking correlation (*r*=0.38 in 42-year-old and *r*=0.47 in 87-year-old). Noteworthy, these results were corroborated in cells without the PIP-FUCCI sensor, to exclude any toxicity effects arising from fluorescence microscopy or viral infection (*r=*0.02 in neonatal, *r=*0.01 in 10-year-old, *r=*0.16 in 21-year-old, *r*=0.49 in 42-year-old and *r=*0.51 in 87-year-old) (Fig. S1D).

Altogether, the results showed that increased duration of all cell cycle phases accounts for cell cycle slowdown along aging, and that mitosis is not temporally insulated from variability in upstream cell cycle events in older cells.

### Defective switch-like activation of CDK1 at mitotic entry in older cells

Since we found mitosis duration to increase with aging and to become coupled with cell cycle length, we asked if the positive feedback regulating the switch-like activation of CDK1 at mitotic entry is compromised in older cells. To test this, we resorted to neonatal and 87-year-old HDFs stably expressing a Cyclin B1-YFP biosensor. Cyclin B1 translocation to the nucleus was used as a proxy of CDK1 activation, since its nuclear translocation depends on direct phosphorylation by CDK1 (Fig. 2A) ^10,18,31,32^. The duration of mitosis was defined as the time between Cyclin B1 nuclear import and its degradation, and as shown previously, it was short and constant in young cells (neonatal HDFs: 28.2 ± 5.5 min, Coefficient of Variation – cv=19.7%) and longer and more variable in old cells (87-year-old HDFs: 49.7 ± 26.37 min, cv=53.1%) (Fig. 2B). To quantitatively measure CDK1 activation, time courses of Cyclin B1 redistribution in individual cells were fitted to the logistic equation, and rise times were calculated from the curve fits for all cells. As expected, CDK1 activation in young cells was illustrated by a sharp sigmoidal curve, typical of an abrupt activation, with a fitted rise time of 2.5 ± 1.1 min (Fig. 2C,D). However, in elderly cells, CDK1 was activated 2.5-fold slower, with a rise time of 6.0 ± 5.1 min, supporting the idea that CDK1 is activated in a more sluggish way (Fig. 2C,D and Fig. S2A).

**Figure 2.**
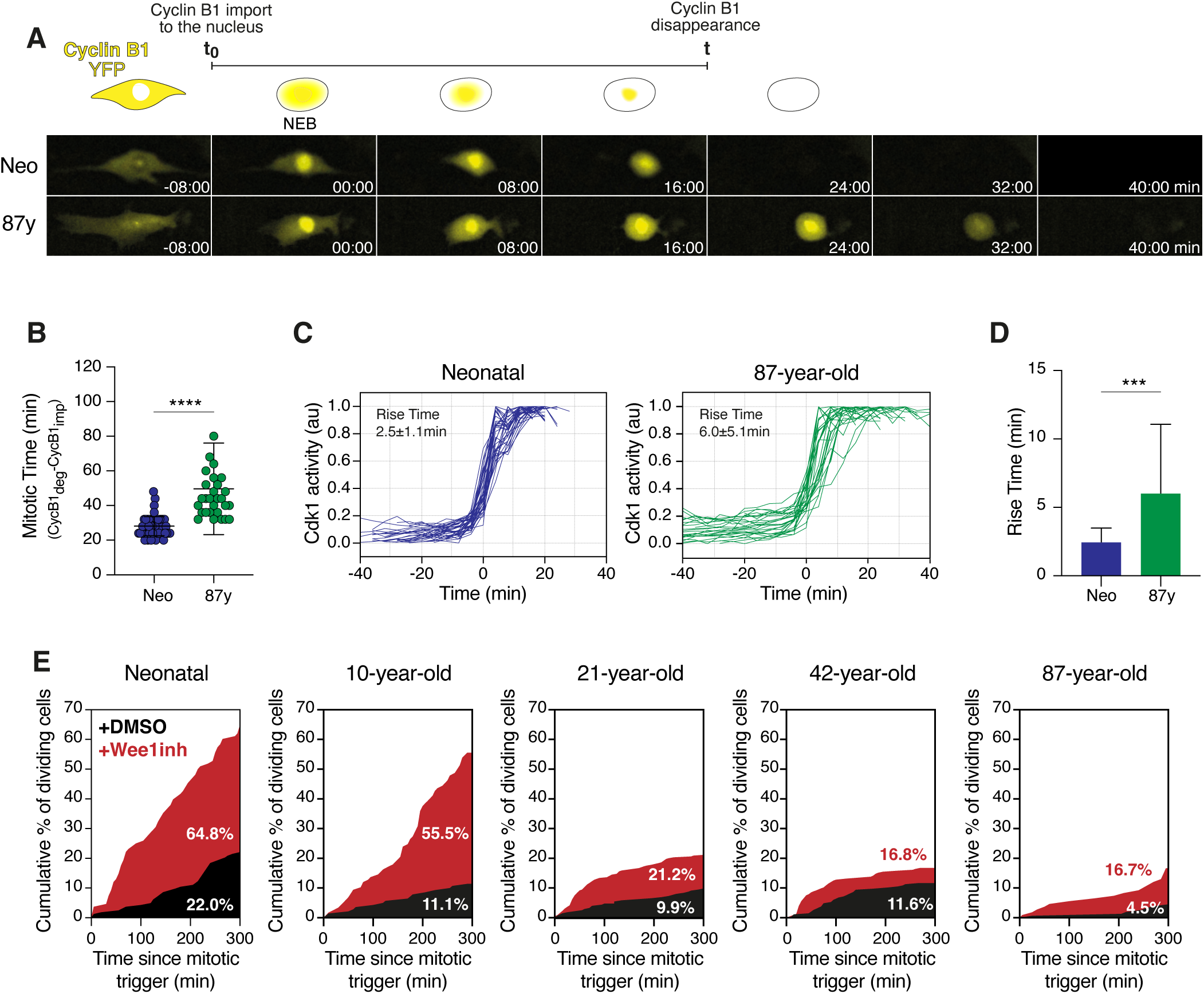
Switch-like CDK1 activation is compromised in older cells. ***A.*** Top: schematic of the appearance of the Cyclin B1 biosensor during mitosis. Bottom: Representative images showing nuclear translocation of Cyclin B1 as a proxy for CDK1 activity in neonatal (top panel) and 87-year-old (bottom panel) cells. ***B.*** Duration of mitosis measured by the time between Cyclin B1 import and Cyclin B1 degradation in neonatal and 87-year-old HDFs. Mean ± SD are shown as bars. **** *p*≤≤0.0001 by Mann-Whitney U test. ***C.*** Quantification of CDK1 activity over time in neonatal (left panel) and 87-year-old (right panel) HDFs. Individual time courses were fitted to the logistic equation y=a+b/1+e^-(t-t0/τ)^ and were scaled to their fitted maximum and minimum values (b and a, respectively) and half-maximal times (t_0_). Rise times (τ) are expressed as mean ± SD (n≥20 cells). ***D.*** Rise times were calculated from the curve fits for single neonatal and 87-year-old-cells represented in C. Mean ± SD are shown as bars. *** *p*≤0.001 by Mann-Whitney U test. ***E.*** Cumulative percentage of dividing cells as a function of time following treatment with mitotic trigger in neonatal, 10-, 21-, 42- and 87-year-old HDFs. Cells were either treated with DMSO (black) or Wee1 inhibitor (Wee1 inh) (red) (n>100 cells, 2 independent experiments).

We next investigated how prone are older cells to undergo mitosis. To this end, live-cell imaging of freely cycling HDFs was performed in the absence or the presence of a mitotic trigger, the small molecule PD 166285 (WEE1 inhibitor) that specifically inhibits MYT1/WEE1, releasing its inhibitory action on CDK1 and enabling entry into mitosis^33,34^. Upon addition of either DMSO or WEE1 inhibitor, cells were imaged for five more hours (Fig. 2E). Within five hours upon DMSO treatment, the percentage of cells entering mitosis decreased gradually along aging, from 22% in neonatal cultures to only 4.5% in octogenarian cultures (Fig. 2E). Responsiveness to the mitotic trigger also decreased progressively with aging, with 64.8% of neonatal cells vs. only 16.7% of 87-year-old cells entering mitosis within five hours upon addition of the WEE1 inhibitor. The refractory period in cells that were unable to divide was 6.5±2.4 hours in neonatal vs. 10.6±4.5 hours in octogenarian cells treated with DMSO, and 2.4 ± 1.0 hrs in neonatal vs. 5.8 ± 4.4 hrs in old cells treated with WEE1 inhibitor (Fig. S2B,C).

Overall, the results showed that CDK1 activation at mitotic entry becomes increasingly sluggish with aging, and older cells exhibit an extended refractory period between two consecutive mitoses in line with their reduced proliferative capacity and responsiveness to a mitotic trigger.

### FOXM1 expression is required for the temporal insulation of mitosis from upstream cell cycle events

Since the repression of transcription factor FOXM1 during aging was previously shown to lead to global shutdown of G2/M genes^19^, including major players of the positive feedback controlling CDK1 activation at mitotic entry (Fig. S3A), we reasoned that it likely accounts for the loss of mitotic insularity with aging. siRNA-mediated depletion of FOXM1 in young cells was used to determine if mitosis becomes dependent on cell cycle length (Fig. S3B). Conversely, FOXM1 overexpression via lentiviral infection in old cells was used to determine if mitotic insularity can be rescued (Fig. S3C). As previously reported^19^, siFOXM1 vs. control neonatal cells exhibited a mitotic delay (57.0 ± 10.8 vs. 43.2 ± 6.6 min) and a prolonged cell cycle (25.1±8.2 vs. 18.4 ± 3.5 hrs) (Fig. S3D). The opposite was observed when a constitutively active truncated form of FOXM1 was overexpressed (OEFOXM1) in 87-year-old cells, with both mitotic duration (42.6 ± 5.9 vs. 55.8 ± 17.4 min) and cell cycle length (20.7 ± 3.2 vs 24.2 ± 10.2 hrs) decreasing in comparison to controls (Fig. S3D). Moreover, siRNA-mediated depletion of FOXM1 in neonatal cells recapitulated the coupling between cell cycle length and mitotic duration (*r*=0.54) as observed in old cells (*r*=0.57) (Fig. 3A), while FOXM1 overexpression in 87-year-old cells restored the mitosis uncoupling from cell cycle length (*r*=0.03) as seen in young cells (*r*=0.04) (Fig. 3A). Next, we measured the time between Cyclin B1 import and its degradation in siFOXM1 cells. We found this time to get longer and more variable in comparison to neonatal control (cv=55.3% vs. cv=27.3%) (Fig. 3B), with a rise time of CDK1 activation of 10.9 ± 6.7 minutes vs. 4.1 ± 2.6 minutes in controls (Fig. 3C,D). On the other hand, overexpression of FOXM1 in elderly cells was able to restore a shorter and more constant mitotic time (cv=33.3% vs. cv=48.0% in old cell controls) (Fig. 3E), as well as a faster activation of CDK1 as depicted by a rise time of 4.9 ± 3.4 minutes, very similar to the rise time in young control cells (4.1 ± 2.6 min) (Fig. 3F,G). Moreover, FOXM1 repression in young cells also increased the time the cells need to be in interphase before being able to respond to a mitotic trigger (Fig. 3H,I), while FOXM1 overexpression in elderly cells decreased their refractory period in response to the WEE1 inhibitor (Fig. 3J,K).

**Figure 3.**
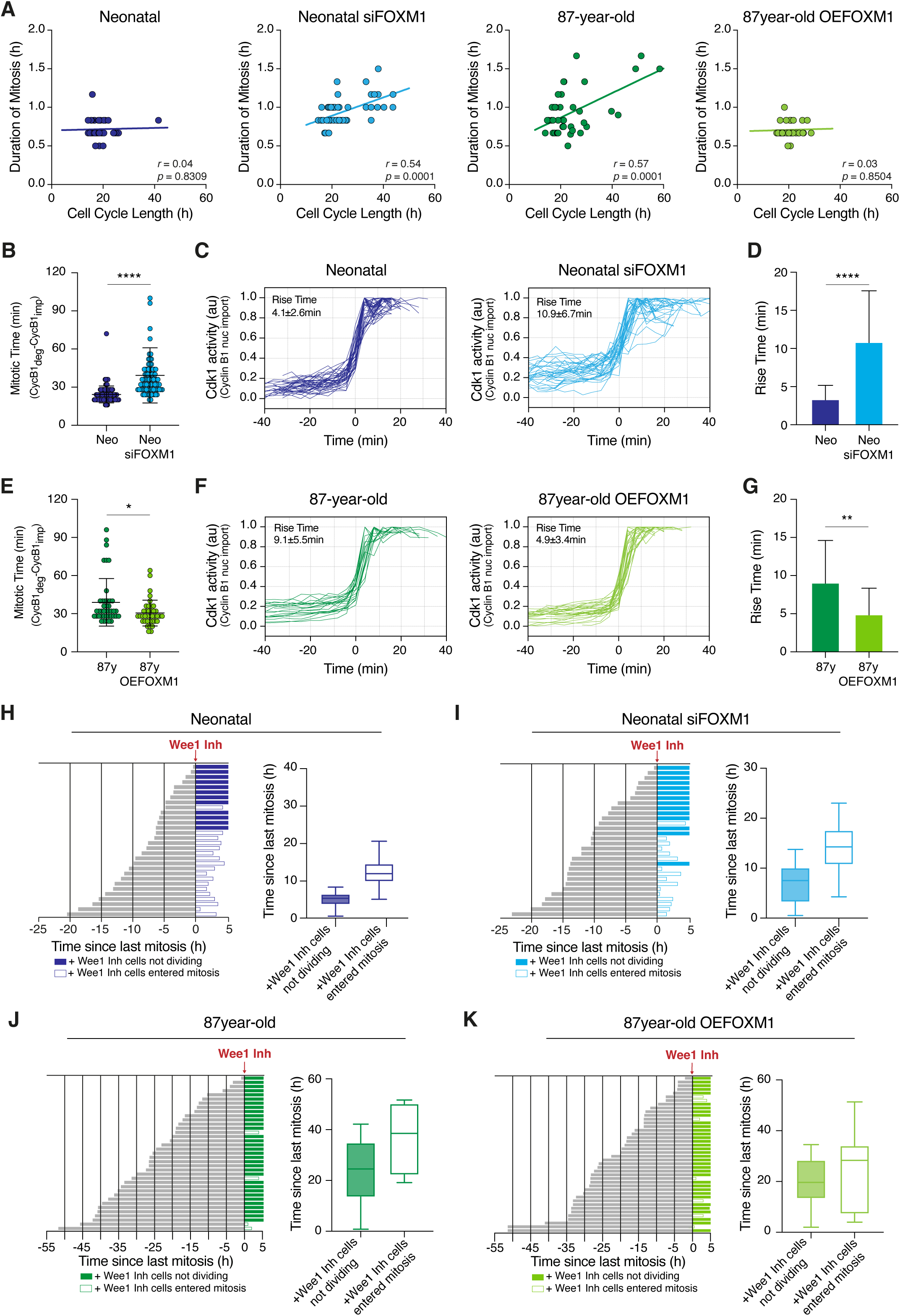
*FOXM1* repression in young cells phenocopies the coupling between cell cycle length and mitotic duration observed in aged cells. ***A.*** Duration of mitosis measured by the time between nuclear envelope breakdown and nuclear envelope reformation as a function of cell cycle length in neonatal HDFs, siFOXM1-depleted neonatal HDFs, 87-year-old HDFs and 87-year-old HDFs overexpressing FOXM1 (OEFOXM1). The trend lines, Pearson coefficient (*r*) and p-values are shown. n>50 cells, ≥2 independent experiments. ***B.*** Duration of mitosis measured by the time between Cyclin B1 import and Cyclin B1 degradation in neonatal and siFOXM1 neonatal HDFs. Mean ± SD are shown as bars. **** *p*≤0.0001 by Mann-Whitney U test. ***C.*** Quantification of CDK1 activity over time in neonatal and siFOXM1 neonatal HDFs. Individual time courses were fitted to the logistic equation y=a+b/1+e^-(t-t0/τ)^ and were scaled to their fitted maximum and minimum values (b and a, respectively) and half-maximal times (t_0_). Rise times (τ) are expressed as mean ± SD (n>25 cells). ***D.*** Rise times were calculated from the curve fits for single neonatal and siFOXM1 neonatal HDFs represented in C. Mean ± SD are shown as bars. **** *p*≤0.0001 by Mann-Whitney U test. ***E.*** Duration of mitosis in 87-year-old and 87-year-old overexpressing-FOXM1 HDFs. Mean ± SD are shown as bars. * *p*≤0.05 by Mann-Whitney U test. ***F.*** Quantification of CDK1 activity over time in 87-year-old and 87-year-old OEFOXM1 HDFs. Rise times (τ) are expressed as mean ± SD (≥14 cells). ***G.*** Rise times were calculated from the curve fits for single 87-year-old and 87-year-old OEFOXM1 HDFs. Mean ± SD are shown as bars. ** *p*≤0.01 by Mann-Whitney U test. ***H.*** Bar plots of neonatal control HDFs representing the time since last division (grey bars) until treated with Wee1 inhibitor (dark blue bars). Filled bars represent no entering into mitosis within 5 hours of treatment with Wee1 inhibitor. Open bars represent cells that entered mitosis (end of the bars represents the time of mitotic entry). Box plot representing the time since last division of neonatal control cells that divided and that did not divide upon treatment with Wee1 inhibitor. ≥2 independent experiments, >60 cells. ***I.*** Bar plots of siFOXM1 neonatal HDFs representing the time since last division (grey bars) until treated with Wee1 inhibitor (light blue bars). Filled bars represent no entering into mitosis within 5 hours of treatment with Wee1 inhibitor. Open bars represent cells that entered mitosis (end of the bars represents the time of mitotic entry). Box plot representing the time since last division of siFOXM1 neonatal HDFs that divided and that did not divide upon treatment with Wee1 inhibitor. ≥2 independent experiments, >100 cells. ***J.*** Bar plots of 87-year-old control (OEFOXM1 -Dox) HDFs representing the time since last division (grey bars) until treated with Wee1 inhibitor (dark green bars). Filled bars represent no entering into mitosis within 5 hours of treatment with Wee1 inhibitor. Open bars represent cells that entered mitosis. Box plot representing the time since last division of 87-year-old control (OEFOXM1 -Dox) that divided and that did not divide upon treatment with Wee1 inhibitor. ≥2 independent experiments, >35 cells. ***K.*** Bar plots of 87-year-old OEFOXM1 (OEFOXM1 +Dox) HDFs representing the time since last division (grey bars) until treated with Wee1 inhibitor (light green bars). Filled bars represent no entering into mitosis within 5 hours of treatment with Wee1 inhibitor. Open bars represent cells that entered mitosis. Box plot representing the time since last division of 87-year-old OEFOXM1 (OEFOXM1 +Dox) HDFs that divided and that did not divide upon treatment with Wee1 inhibitor. ≥2 independent experiments, >35 cells.

Taken together, the data showed that FOXM1 repression in young cells phenocopies the changes in cell cycle dynamics observed in aged cells. Contrarywise, FOXM1 overexpression in aged cells rescues the CDK1 switch-like activation required for the temporal insulation of mitosis from upstream cell cycle events and improves the proliferative capacity of aged cells by decreasing the refractory period between two consecutive mitoses.

### Increased FZR1/Cdh1 levels lead to FOXM1 repression and loss of proliferative capacity in aged cells

Proteolytic degradation of FOXM1 at mitotic exit is regulated by the APC/C^Cdh1^ E3-ubiquitin ligase activity, so that when APC/C^Cdh1^ is on, FOXM1 activity is off, and vice-versa (Fig. 4A)^23^. Thus, we asked if increased FZR1/Cdh1 levels correlate with FOXM1 repression during aging. Indeed, we found Cdh1 levels to gradually increase in fibroblasts retrieved from advancing age donors, and concomitantly to the steady decrease of FOXM1 levels (Fig. 4B and S4A). In addition, siFOXM1 depletion in neonatal cells was found to increase Cdh1 levels (Fig. 4C and S4B). These results, together with the fact that FOXM1 is activated by CDK1-Cyclin B1 and that APC/C^Cdh1^ drives both Cyclin B1 and FOXM1 proteolytic degradation, suggest for a feedback regulation axis between CDK1, FOXM1 and APC/C^Cdh1^ that appears to primarily account for age-related changes in cell cycle dynamics. We therefore went on assessing if modulation of Cdh1 levels, alike modulation of FOXM1 levels, can impact cell proliferation and senescence.

**Figure 4.**
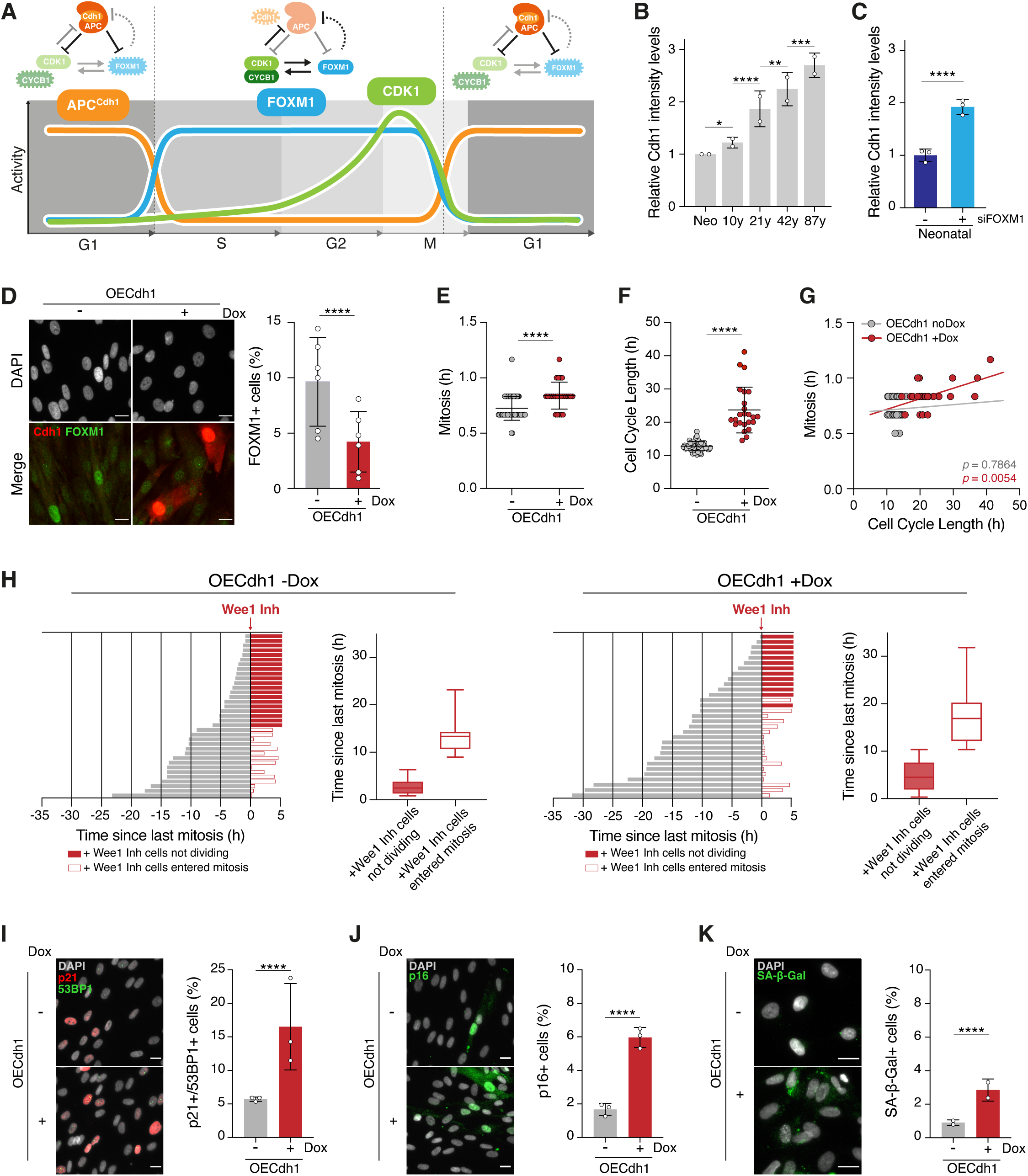
Overexpression of Cdh1 in young cells reproduces aged cells’ phenotypes. ***A.*** Feedback regulation between CDK1, FOXM1 and APC/C^Cdh1^ controls cell cycle dynamics. ***B.*** Quantification of Cdh1 intensity levels by immunofluorescence analysis of neonatal, 10-, 21-, 42- and 87-year-old HDFs. Levels were normalized to neonatal. n≥3250 cells, 2 independent experiments. All values represent mean ± SD. * *p*≤0.05, ** *p*≤0.01, *** *p*≤≤0.001, **** *p*≤0.0001 by two-sided Kruskal-Wallis with Dunn’s multiple comparison correction. ***C.*** Quantification of Cdh1 intensity levels by immunofluorescence analysis of mock- and siFOXM1-depleted neonatal HDFs. Levels were normalized to mock. n≥3330 cells, 3 independent experiments. All values represent mean ± SD. **** *p*≤0.0001 by Mann-Whitney U test. ***D.*** Representative images and quantification of FOXM1-positive cells by immunofluorescence analysis in neonatal control (OECdh1 -Dox) or overexpressing Cdh1 (OECdh1 +Dox) HDFs. Levels were normalized to neonatal control. n≥1100 cells, 6 independent experiments. All values represent mean ± SD. **** *p*≤0.0001 by two-tailed Fisher’s exact test. Scale bar, 20μm. ***E-F.*** Mitotic duration (E) and cell cycle length (F) measured in neonatal control (OECdh1 -Dox) or overexpressing Cdh1 (OECdh1 +Dox) cells. n>50 cells, 2 independent experiments. All values represent mean ± SD. **** *p*≤0.0001 by Mann-Whitney U test. ***G.*** Duration of mitosis measured by the time between NEB and NER as a function of cell cycle length in neonatal control (OECdh1 -Dox) or overexpressing Cdh1 (OECdh1 +Dox) cells. The trend lines and p-values are shown. n>50 cells, ≥2 independent experiments. ***H.*** Bar plots of neonatal cells, control (OECdh1 -Dox) or overexpressing Cdh1 (OECdh1 +Dox) as indicated, representing the time since last division (grey bars) until treated with Wee1 inhibitor (red bars). Filled red bars represent no entering into mitosis within 5 hours of treatment with Wee1 inhibitor. Open red bars represent cells that entered mitosis (end of the bars represents the time of mitotic entry). Box plots representing the time since last division of neonatal control or overexpressing Cdh1 HDFs treated with Wee1 inhibitor that did not divide (filled bar) and that divided (open bars) in response to Wee1 inhibitor (n>30 cells). ***I.*** Representative images and quantification of neonatal control or overexpressing Cdh1 cells staining positive for both p21/CDKN1A (cell cycle inhibitor) and 53BP1 (≥ 1 foci; DNA damage) senescence markers. All values represent mean ± SD. n≥3000 cells, 3 independent experiments. **** *p*≤0.0001 by two-tailed Fisher’s exact test. Scale bar, 20μm. ***J.*** Representative images and quantification of neonatal control or overexpressing Cdh1 cells staining positive for p16/CDKN2A senescence marker. All values represent mean ± SD. n≥3000 cells, 3 independent experiments. **** *p*≤0.0001 by two-tailed Fisher’s exact test. Scale bar, 20μm. ***I.*** Representative images and quantification of neonatal control or overexpressing Cdh1 cells staining positive for SA-β-gal activity assay. All values represent mean ± SD. n≥3000 cells, 2 independent experiments. **** *p*≤0.0001 by two-tailed Fisher’s exact test. Scale bar, 20μm.

Lentiviral transduction of young cells with a doxycycline (dox)-inducible transgene was used to overexpress FZR1/Cdh1 (OECdh1). Cdh1 levels were 5-fold higher in transduced young cells (Fig. S4C and S4D), resulting in decreased FOXM1 levels (Fig. 4D and S4E). As expected for FOXM1 repression, both cell cycle and mitotic durations were increased in young cells overexpressing Cdh1 in comparison to control cells (cell cycle duration, 23.7 ± 6.9 vs. 12.8 ± 1.4 hrs; mitosis duration, 50.4 ± 7.2 vs. 43.8 ± 6.6 min) (Fig. 4E,F). Moreover, Cdh1 overexpression was able to disrupt the temporal insulation of mitosis which became correlated with total cell cycle length (*r*=0.30 vs. *r*=0.04 in control cells). Also, the proliferative capacity of young cells overexpressing Cdh1 was assessed, and like older cells, they became poorly responsive to WEE1 inhibitor, exhibiting an extended lag time in interphase before commitment into a new round of mitosis (Fig. 4H).

Finally, considering that FOXM1 repression induces senescence, we measured the levels of senescence markers in OECdh1 cells. The percentage of cells exhibiting markers of DNA damage, cell cycle inhibitors and senescence-associated β-galactosidase activity increased in OECdh1 vs. control cells (Fig. 4I-K). In agreement with senescence accrual, both mitotic and proliferative indexes were decreased in OECdh1 cell cultures (Fig. S4F,G). Overall, the results showed that increased levels of Cdh1 are sufficient to impact cell cycle dynamics leading to loss of proliferative capacity and senescence accrual.

### FZR1/Cdh1 repression in elderly cells restores cell cycle dynamics and rescues senescence

Next, we investigated if younger phenotypes could be rescued in elderly cells by conversely downregulating Cdh1 expression. siRNA-mediated depletion of Cdh1 (80% knockdown) in 87-year-old cells (Fig. 5A and Fig. S5A,B) was sufficient to translate into a 2-fold increase in FOXM1 levels in comparison to controls (Fig. S5C). Accordingly, mitotic duration and cell cycle length decreased in siCdh1 vs. control cells (mitosis: 41.4 ± 7.2 vs. 45.6 ± 9.6 min; cell cycle: 16.2 ± 4.4 vs. 22.2 ± 6.9 hrs) (Fig. 5B,C). Also, Cdh1 knockdown in old-age cells restored mitosis insularity from the cell cycle (*r*= 0.007 vs. *r*=0.65 in controls) (Fig. 5D), as well as the responsiveness to the WEE1 inhibitor mitotic trigger (Fig. 5E and Fig. S5D). Consistently to an improved cell cycle dynamics in siCdh1-depleted elderly cells, we found increased mitotic and proliferative indexes (Fig. S5E,F) and decreased levels of senescence markers (Fig. 5F-H). Altogether, the results showed that Cdh1 repression acts to restore cell cycle dynamics in aged cells while preventing senescence accrual.

**Figure 5.**
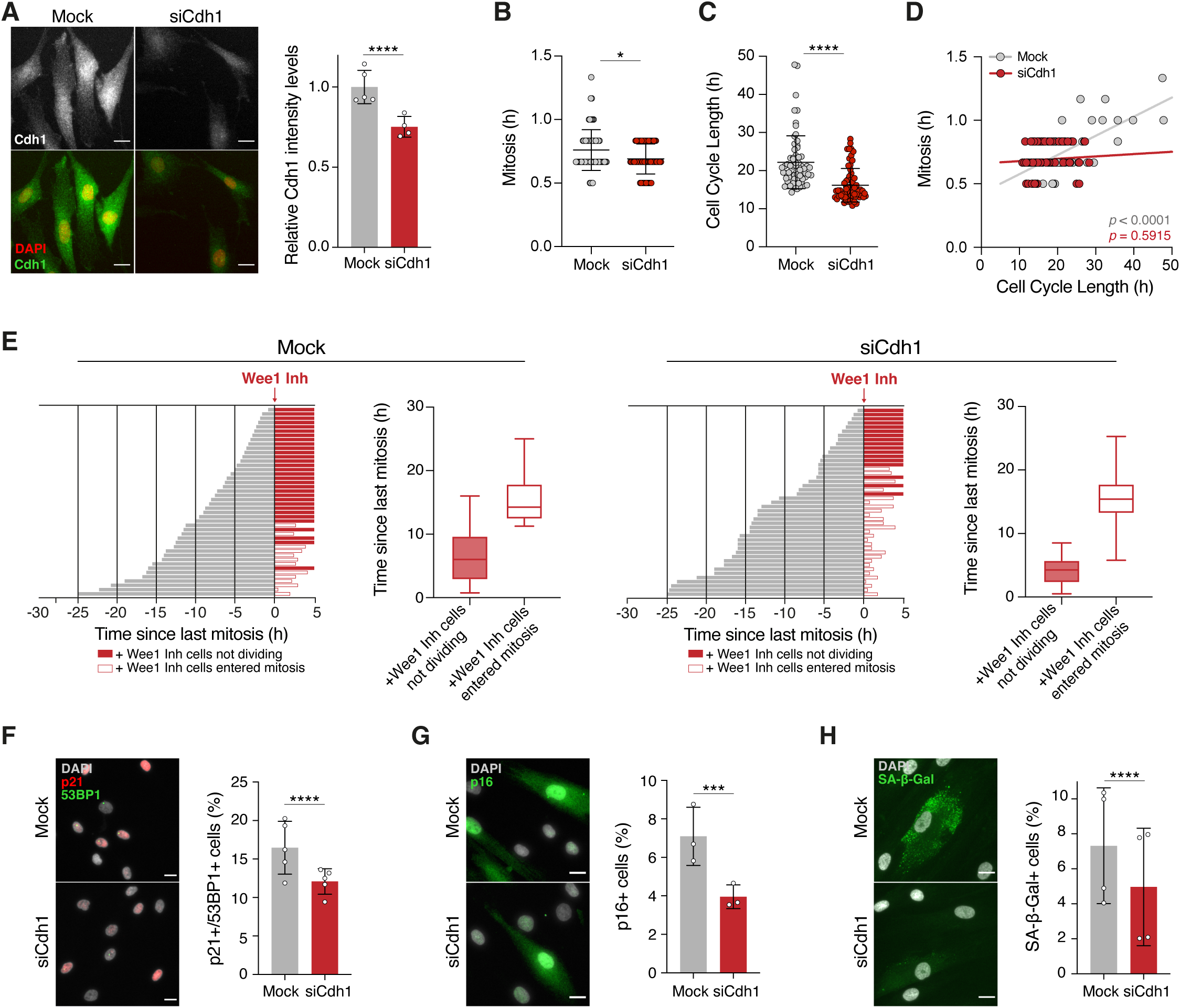
*FZR1/Cdh1* repression in elderly cells restores cell cycle dynamics and rescues senescence. ***A.*** Representative images and quantification of Cdh1 intensity levels by immunofluorescence analysis in mock- and siCdh1-depleted aged HDFs. Levels were normalized to mock control. All values represent mean ± SD. n≥330 cells, ≥4 independent experiments, 2 biological replicates. *\*\*\*\* p*≤0.0001 by Mann-Whitney U test. Scale bar, 20μm. ***B-C.*** Mitotic duration (time between NEB and NER) (C) and cell cycle length (D) measured in mock- and siCdh1-depleted 87-year-old HDFs. All values represent mean ± SD. n>60 cells, 2 independent experiments. ***D.*** Duration of mitosis as a function of cell cycle length in mock and siCdh1-depleted aged cells. The trend lines and *p-values* are shown. n>60 cells, ≥2 independent experiments. ***E.*** Bar plots of mock or siCdh1-depleted 87y-year-old HDFs representing the time since last division (grey bars) until treated with Wee1 inhibitor (red bars). Filled red bars represent no entering into mitosis within 5 hours of treatment with Wee1 inhibitor. Open red bars represent cells that entered mitosis (end of the bars represents the time of mitotic entry). Box plots representing the time since last division of mock- or siCdh1-depleted 87y-year-old HDFs treated with Wee1 inhibitor that did not divide (filled bar) and that divided (open bar) in response to Wee1 inhibitor (n>65 cells). ***F.*** Representative images and quantification of mock or siCdh1-depleted 87y-year-old HDFs staining positive for both p21/CDKN1A and 53BP1 (≥ 1 foci) senescence markers. All values represent mean ± SD. n≥2500 cells, 5 independent experiments, 2 biological replicates. **** *p*≤0.0001 by two-tailed Fisher’s exact test. Scale bar, 20μm. ***G.*** Representative images and quantification of mock or siCdh1-depleted 87y-year-old HDFs staining positive for p16/CDKN2A senescence marker. All values represent mean ± SD. n≥1500 cells, 3 independent experiments. *** *p*≤≤0.001 by two-tailed Fisher’s exact test. Scale bar, 20μm. ***H.*** Representative images and quantification of mock or siCdh1-depleted 87y-year-old HDFs staining positive for SA-β-gal activity assay. All values represent mean ± SD. n≥2000 cells, 4 independent experiments. **** *p*≤0.0001 by two-tailed Fisher’s exact test. Scale bar, 20μm.

### CDK1:FOXM1:APC/C^Cdh1^ feedback regulation in a mother cell imprints the proliferative capacity of its daughter cells

Lastly, and based on previous studies showing that prolonged mitosis of a mother cell impacts daughter cells’ fate^18,35,36^, we investigated if changes in the CDK1:FOXM1:APC/C^Cdh1^ feedback loop impinge memories on the next cell cycle. Asynchronous cultures of young, old, and siCdh1-depleted old HDFs were imaged by long-term time-lapse microscopy to measure mitotic duration of mother cells, and the levels of cell cycle inhibitors (*CDKN1A*/p21 and *CDKN2A*/p16) were assessed afterwards in daughter cells to ascertain the effect of a mitotic delay on the progeny proliferative capacity (Fig. 6A). As abovementioned, mitosis is short and constant in young cells (34.8 ± 5.0 min), longer and variable in older cells (44.1 ± 8.9 min), turning short and constant in the later if Cdh1 is depleted (37.2 ± 5.8 min) (Fig. 6B). Using live-cell fixed-cell correlative microscopy analysis, we found that, in young cells, the mitotic duration of the mother cell correlates with the accumulation of p21 levels in the daughter cells (*r*=0.17), suggesting that even minor (<20 min) mitotic delays allow the assembly of the ‘mitotic stopwatch’ complex comprised by USP28-53BP1-p53 and reported to mediate CDKN1A/p21 accrual in the next cell division cycle ^36^ (Fig. 6C and Fig. S6A). Intriguingly, although old cells exhibit higher (>20 min) mitotic delays, a correlation between mitotic duration of mother cells and p21 levels in their daughter cells was not observed (*r*=0.078). Independently of the mother cells’ mitotic duration, daughter cells invariably exhibited 2-fold higher p21 levels than young counterparts, indicating that aged mother cells are already primed with higher p21 levels before mitosis (Fig. 6C and S6A). Indeed, *CDKN1A/p21* transcript levels were found to be higher in mitotic cells isolated from aged vs. young HDF cultures^19^. Moreover, transcript levels of *USP28*, as well as of *PLK1,* which is required for the assembly of the USP28-53BP1-p53 complex, were found to be downregulated in aged vs. young mitotic HDFs (Fig. S6A), compliantly with an apparently inoperative ‘mitotic stopwatch’ during aged cell mitosis. Also, daughter cells from older mothers did not accumulate p21 levels the longer they spent in interphase (r=0.017) as seen for young counterparts (r=0.20) (Fig. S6B). Importantly, and demonstrating the effect of CDK1:FOXM1:APC/C^Cdh1^ feedback regulation in mitotic delay memory, FZR/Cdh1 depletion in older cells was able to restore the correlation between mitotic duration of mother cells and p21 levels in daughter cells as seen for young controls (Fig. 6C and S6B).

**Figure 6.**
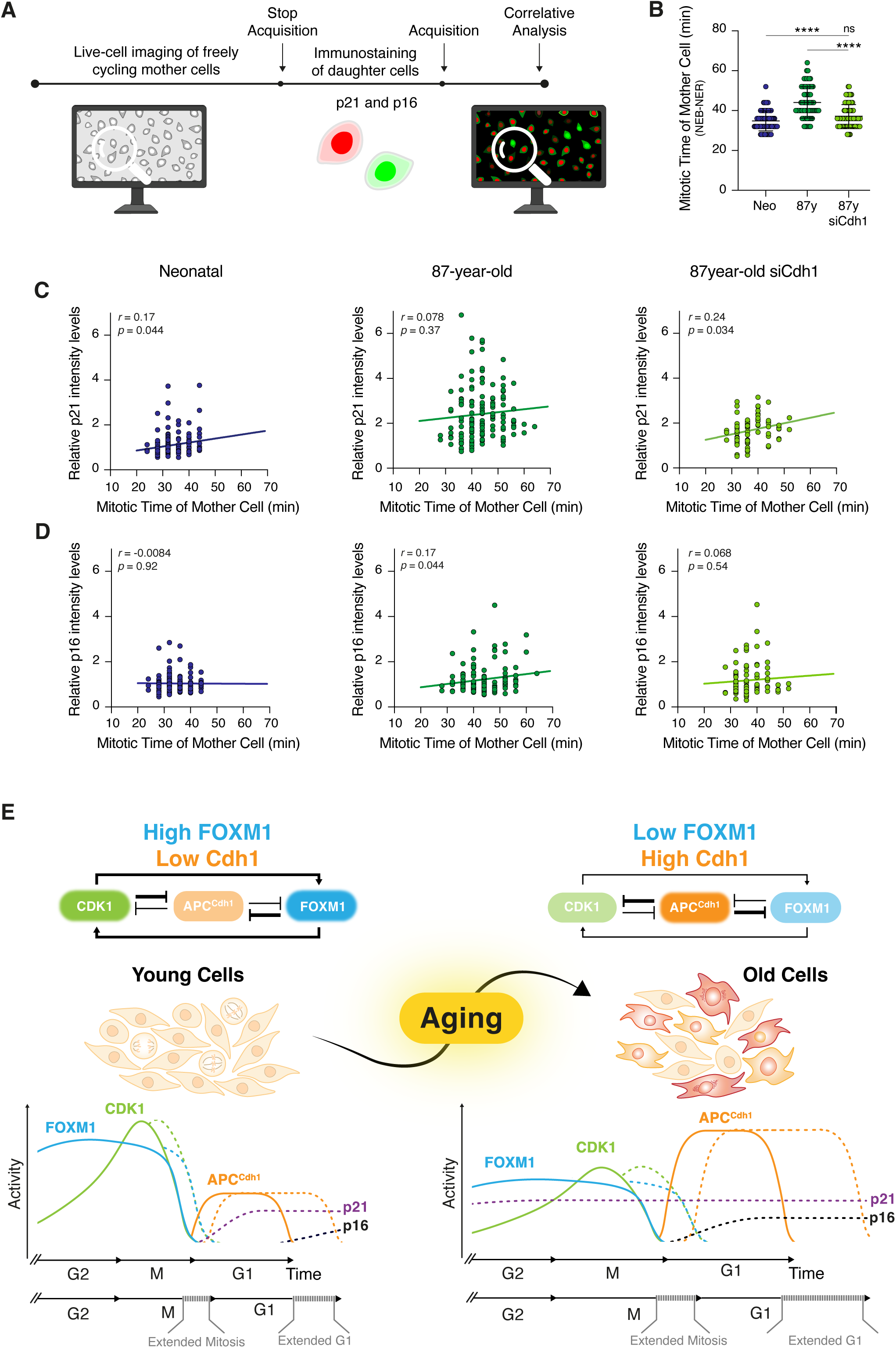
Lengthened mitoses of aged mother cells imprint their progeny with increased levels of p16 cell cycle inhibitor. ***A.*** Schematic representation of the live cell-fixed cell correlative microscopy analysis. ***B.*** Duration of mitosis measured by the time between nuclear envelop break down (NEB) and nuclear envelope reformation (NER) in neonatal, 87-year-old and siCdh1-depleted 87y-year-old HDFs. Mean ± SD are shown as bars. ns, non-significant and **** *p*≤0.0001 by Mann-Whitney U test. ***C.*** Correlation between relative p21 intensity levels of daughter cells and the mitotic time of mother cells in neonatal, 87-year-old and siCdh1-depleted 87-year-old cultures. The trend lines, Pearson coefficient (*r*) and p-values are shown. n>60 cells. ***D.*** Correlation between relative p16 intensity levels of daughter cells and the mitotic time of mother cells in neonatal, 87-year-old and siCdh1-depleted 87y-year-old cultures. The trend lines, Pearson coefficient (*r*) and p-values are shown. n>60 cells. ***E.*** Schematic summary of the CDK1:FOXM1:APC/C^Cdh1^ feedback regulation determining the changes in cell cycle dynamics during aging. Young cells with high CDK1:FOXM1 activities only accumulate CDKN1A/p21 if there is a mitotic delay, which then leads to extended G1 and CDKN2A/p16 upregulation in the daughter cells. Aged mitotic cells, already primed with higher levels of CDKN1A/p21, exhibit extended mitotic duration due to low CDK1:FOXM1 activities, which in turn impart a G1 delay in the next cell cycle due to upregulation of APC/C^Cdh1^ and CDKN2A/p16 activities.

We additionally measured CDKN2A/p16 levels in the daughter cells, another cell cycle inhibitor operating at a later stage of the senescence state^37^. Whereas p16 levels were low in the daughter cells from young mothers and did not increase with mitotic delay (r=-0.084), in the daughter cells from older mothers we found p16 levels to increase with mitotic duration (*r*=0.17), a correlation that was lost upon Cdh1 depletion (r=0.068) (Fig. 6D). This indicates that a prolonged mitosis in aged mother cells with higher p21 levels imprints their daughter cells for higher p16 levels. Accordingly, p16 levels in daughter cells from older mother cells did not correlate with the time the daughter cells spent in interphase (r=0.059), in contrast to the positive correlation found in daughter cells from younger or siCdh1-depleted older mother cells (r=0.26 and r=0.21, respectively) (Fig. S6C).

Taken together the results showed that whereas in young mother cells an increased mitotic duration imprints daughter cells for higher p21 (but not p16) levels, in the daughter cells from older mothers p21 levels are invariably 2-fold higher, and the mitotic delay alternatively imprints daughters for higher p16 levels and, consequently, propensity for a senescence outcome (Fig. 6E). Importantly, Cdh1 repression in older dividing cells, rescues the p16 levels imprinted by the mothers’ mitotic delay in the daughter cells, which similarly to daughter cells from young mothers, rather accumulate p21 and p16 levels the longer they spend in interphase.

## DISCUSSION

Aging is characterized by impaired tissue regeneration, that eventually leads to organ failure. One contributing factor is the loss of proliferative capacity during aging. Here, using a PIP-FUCCI sensor^30^, we demonstrated that all cell cycle phases are prolonged as the cells age, with G1-phase extending the most. Thus, alike G1-phase lengthening during early development to allow for cell commitment to differentiation^38–40^, and similarly to extended cell cycle gap phases in somatic vs. embryonic cells to allow for checkpoint control of genomic integrity ^7^, also aged cells remodel and lengthen their cell division cycle to possibly cope with increased molecular damage ^41^.

As an exemption to previous findings showing that mitosis is invariably short and constant regardless of upstream cell cycle events ^18^, we found that mitotic duration gets variable and coupled with cell cycle length along aging due to weakened positive feedback controlling CDK1-CyclinB1 activation. Defective CDK1-CyclinB1 activation at mitotic entry in old cells leads to sluggish mitotic progression in line with our previous findings showing increased mitotic defects and activation of the mitotic checkpoint ^19^. Moreover, in agreement with the quantitative model for orderly and unidirectional cell cycle progression^42–44^, and indicating that elder cells take longer to reach the threshold levels of CDK activities needed for cell cycle phase transitions, we found elder cells to exhibit weak responsiveness to a mitotic trigger and to spend more time in interphase before entering mitosis.

Furthermore, we disclosed that high levels of the FOXM1 transcription factor are required to keep mitosis short, constant, and temporally insulated from upstream cell cycle events, since FOXM1 drives the transcription of all molecular players of the positive feedback loop, thereby determining its strength^19,45–48^. We previously demonstrated that *FOXM1* repression is the major driver of age-associated mitotic decline. Here, we show that low FOXM1 levels in aged cells, or experimentally induced (siRNA) in young cells, lead to defective CDK1 activity, loss of mitotic insularity and weak responsiveness to a mitotic trigger. Low CDK1 activity in turn further impairs FOXM1 activation and transcription of Cyclin B1 ^20^. Importantly, all the changes in cell cycle dynamics were restored in aged cells upon overexpression of FOXM1. Cell cycle boosting might therefore account for the previously reported senotherapeutic effect of an extra copy of FOXM1 in animal models of natural and accelerated aging^25^.

In addition, we showed that both old cells and siFOXM1-depleted young cells accumulate high levels of FZR1/Cdh1, the co-activator of the APC/C E3-ubiquitin ligase driving FOXM1 proteolysis at mitotic exit. Interestingly, *FZR1/Cdh1* is highly expressed in post-mitotic neurons to sustain neurons in a non-dividing state via the ubiquitin-mediated degradation of cell cycle proteins^28,49,50^. This suggests that similarly to neurons, high Cdh1 levels limit the proliferative capacity of aged cells. Indeed, overexpression of *FZR1/Cdh1* in young cells recapitulated phenotypes observed in elder and siFOXM1-depleted cells, namely loss of mitotic insularity and proliferative capacity, as well as accelerated accumulation of senescence markers (p21/53BP1 levels, p16 levels and lysosomal mass) ^51–54^. Critically, *FZR/Cdh1* knockdown in older cells, restored FOXM1 levels, increased proliferative capacity, and delayed senescence. Considering that Cdh1 levels and APC/C^Cdh1^ activity increase during the differentiation of human embryonic stem cells ^27^ ^55^, one can speculate that the cell cycle remodels twice during lifetime: when the cells transit from a pluripotent to a differentiated state ^56^, and later on when cells transit to an aged state.

Finally, we found that the age-associated coupling of mitotic duration with previous cell cycle events also impacts the subsequent cell cycles. Aged cells with prolonged cell cycle and mitotic delay imprint their daughter cells for higher levels of cell cycle inhibitors in G1, and thereby, loss of proliferative capacity. This is in agreement with previous work showing that lengthened mitoses impair the daughter cells from reaching another round of mitosis^18^. However, the accumulation of cell cycle inhibitors in the daughter cells from aged mother cells appears not due to the recently identified mitotic stopwatch surveillance mechanism based on p53 stabilization via assembly of the USP28-53BP1-p53 complex during a mitotic delay^36^. Due to FOXM1 repression, aged mitotic cells have low levels of PLK1^19^, which activity is required to drive the assembly of the mitotic stopwatch complex ^36^. Moreover, the mitotic delay in aged cells is minor in comparison to the delay threshold reported as needed to activate the mitotic stopwatch ^36^. With an apparently inoperative mitotic stopwatch, how do aged mother cells imprint their daughters to cell cycle inhibition? We found that increased FZR1/Cdh1 levels in aged cells are causally linked to the inherited memory of a mitotic delay, since siCdh1 depletion was able to rescue the accrual of cell cycle inhibitors in the daughter cells. Just recently, the p53 ubiquitin ligase MDM2 was reported as a mitotic length timer in the context of stochastic delays in normal cells ^57^. However, we found *MDM2* transcript levels to increase in aged mitotic cells^19^, thereby suggesting that FZR1/Cdh1 ubiquitin ligase activity might rather dictate memory for G1 delay in the next cell cycle in aging. Noteworthy, a correlation between mother cell mitotic duration and p16 accrual was found in the progeny of aged cells, explaining their propensity to a senescent fate in comparison to the progeny of young cells in which p21 accrual instead correlates with the mother cell mitotic delay.

In sum, the mitotic fitness decline during aging traces the proliferative capacity of daughter cells via the CDK1:FOXM1:APC/C^Cdh1^ feedback regulatory loop. These molecular players allow the partition of the cell cycle in two modules – interphase and mitosis – which length and accurate progression impacts one another. Both modules progressively increase as the cells age, explaining why cell cycle slows down, ultimately leading to cell cycle arrest and senescence onset. The imprinting of cell cycle inhibitor levels in new-born cells determines their tempo for cell cycle commitment. Moreover, we show that different cell division cycles can be generated by simply changing the levels of FOXM1 or FZR/Cdh1 feedback players, highlighting the rejuvenating potential of modulating cell cycle fitness ^19,25,29^.

## MATERIALS AND METHODS

### Cell Culture

All cell stocks were routinely tested for mycoplasma and were cultured at 37°C and humidified atmosphere with 5% CO_2_. Human dermal fibroblasts established from skin samples of reported “healthy” Caucasian males with ages ranging from neonatal to octogenarian (neonatal, DFM021711A from ZenBio Biobank; 10-year-old, GM03348; 21-year-old, GM23964; 42-year-old, AG06235; 75-year-old AG11073 and 87-year-old, AG10884; all from Coriell Cell Repository) were cultured in Minimal Essential Medium (MEM) with Earle’s salts and L-glutamine (10-010-CV, Corning, VA, USA), supplemented with 15% Fetal Bovine Serum (FBS) (Gibco, Paisley, UK) and 1x Antibiotic-Antimycotic (Gibco, NY, USA). Only early passage dividing fibroblasts (up to passage 3–5) with cumulative population doubling level PDL<24 was used in all experiments. To produce lentiviruses, HEK293T cells were cultured in Dulbecco’s modified Eagle’s medium (DMEM) (Gibco, Paisley, UK) supplemented with 10% FBS (Gibco, Paisley, UK) and 1x Antibiotic-Antimycotic (Gibco, NY, USA).

### Lentiviral Generation and Biosensors

To generate pLVX–Tight-Puro–FZR1, a *Not* I–*FZR1*–*EcoR* I fragment was amplified from HA-Cdh1 Addgene plasmid (Catalog no.#11596) with primers *Fw: 5’-ATATAGCGGCCGCATGGACCAGGACTATGAGC-3’* and *Rv: 5’-ATATAGAA TTCTTACCGGATCCTGGTGAAG-3’*. The fragment was then inserted into pLVX– Tight-Puro (Clontech) digested with *Not* I + *EcoR* I. HEK293T helper cells were transfected with packaging plasmids pMd2.G and psPAX2 using Lipofectamine 2000 (Life Technologies, CA, USA) to generate lentiviruses carrying pLVX–Tight-Puro– Cdh1, pCSII-EF1a-CyclinB1-YFP ^18^, pLenti-PGK-Neo-PIP-FUCCI ^30^ (Addgene #118616) or pLVX–Tight-Puro–FOXM1-dNdK ^19^. Virus-containing media was collected 48 hrs after transfection and spun down at 2400 rpm for 5 min. The supernatant was aliquoted and snap-frozen using liquid nitrogen before storage at - 80°C. HDFs were infected for 6 hrs in the presence of 8 μg/mL polybrene (AL-118, Sigma-Aldrich, MO, USA). In the case of pLVX–Tight-Puro–Cdh1 and pLVX–Tight-Puro–FOXM1-dNdK, cells were co-infected with the responsive and the reverse tetracycline-controlled transactivator (rTTA) lentiviruses at a ratio 2:1. In the following day, 500ng/mL of doxycycline (D9891, Sigma-Aldrich, MO, USA) was added to the medium to induce co-transduction. Phenotypes were analysed and quantified 72 hrs after induction. Regarding pCSII-EF1a-Cyclin B1-YFP and pLenti-PGK-Neo-PIP-FUCCI cells were sorted one week after infection in a FACS Aria II cell sorter (BD Biosciences, CA, USA) using an 85μm nozzle and the blue (488nm) and yellow/green (561nm) lasers. Cells were gated by forward scatter area (FSC-A) vs. side scatter area (SSA-A) and FSC-A vs. FSC-height (FCS-H) plots to exclude dead cells and doublets/clumps, respectively. The sorting gates were designed accordingly to the respective auto-fluorescent control.

### siRNA knockdown

Cells were plated in serum-depleted culture medium and, after 1 hr, transfected either with 50nM of a small interfering RNA (siRNA) against *FOXM1* (SASI_Hs01_00243977 from Sigma-Aldrich, MO, USA), or with 25nM of siRNA against *FZR1*/Cdh1 (M-015377-01 from Dharmacon, CO, USA). Transfections were performed using Lipofectamine RNAiMAX (Invitrogen, CA, USA) in Opti-MEM medium (Gibco, Paisley, UK) according to the manufacturer’s instructions. Transfection medium was replaced by complete medium after 6 hrs. Phenotypes were analysed and quantified 72 hrs after transfection for siRNA against *FOXM1* and 48 hrs for siRNA against *FZR1*/Cdh1.

### Immunofluorescence and acquisition

Cells were seeded 2-3 days before fixation in 96-well plates (#89626 from Ibidi GmbH, Gra□felfing, Germany). Cells were rinsed twice in Phosphate buffered saline (PBS) before fixation with 4% paraformaldehyde in PBS for 10 min at room temperature. Cells were washed 3 times for 5 min with PBS before permeabilization in PBS with 0.3% Triton X-100 at room temperature for 7 min. Cells were then washed 3 times with PBS-T (PBS + 0.05% Tween-20) and incubated with blocking buffer containing 10% FBS in PBS-T for 1 hr at room temperature. Cells were incubated overnight at 4°C with primary antibodies diluted in PBS-T + 5% FBS as follows: mouse anti-FZR1 (#SC-56312, Santa Cruz Biotechnology, CA, USA) 1:75, rabbit anti-53BP1 (#4937, Cell Signalling Technology, MA, USA), 1:500; mouse anti-p21/CDKN1A (#SC-6246, Santa Cruz Biotechnology, CA, USA), 1:800; rabbit anti-FOXM1 (#13147, ProteinTech Group, Inc., IL, USA), 1:500; mouse anti-KI67 (8D5, #9449, Cell Signalling Technology, MA, USA), 1:500; rabbit anti-p16^INK4a^/CDKN2A (#ab7962, Abcam, Cambridge, UK), 1:200; rabbit anti-phospho-H3 (#06-570, Millipore, Billerica, MA), 1:1500. After being washed 3 times with PBS-T, cells were incubated with secondary antibodies Alexa Fluor-488 and/or Alexa Fluor-568 (Life Technologies, CA, USA) diluted 1:1000 in PBS-T + 5% FBS for 1 hr at room temperature. Nuclei were stained with 1µg/mL DAPI (Sigma-Aldrich, MO, USA).

### SA-**β**-gal activity assay

Live cells were incubated for 90 min in medium with 100 nM Bafilomycin A1 (B1793, Sigma-Aldrich, MO, USA) to induce lysosomal alkalinization. Fluorogenic substrate for β-galactosidase, fluorescein di-β-D-galactopyranoside (F2756, Sigma-Aldrich, MO, USA), was then added to the medium (33 μM), and incubation carried out for another 90 min. Cells were then fixed in 4% paraformaldehyde for 15 min, rinsed with PBS once and permeabilized with 0.1% Triton-X100 in PBS for 15 min. Nuclei were stained with 1µg/mL DAPI (Sigma-Aldrich, MO, USA).

### Fixed-cell imaging

For fixed-cell immunostaining analyses, cells were imaged in the inverted motorized Leica DMI6000B widefield microscope (Leica Microsystems) equipped with Hamamatsu Orca-FLASH4.0 sCMOS (C11440, Hamamatsu) and Leica EL6000 light source (metal-halide lamp), and controlled through the LAS X software (version 3.8.1.26810). Samples were imaged using a binning 2×2. DAPI acquisition was done using the AT filter cube set (ex 340-380 - bs 400 - em LP 425), with FIM at 100%, and an exposure time of 20ms. For Alexa-488, the L5 filter cube set (ex 460/40m - bs 505 - em 527/30) was used, with FIM at 100% and an exposure time of 1s. For Alexa-568, the TX2 filter cube set (ex 560/40 - bs 660 - em 700/75) was used, with FIM at 100% and an exposure time of 1s.

### Time-lapse live-cell imaging

For live-cell imaging experiments, cells were seeded in 96-well or 24-well plates (#89626 and #82426 from Ibidi GmbH, Gra□felfing, Germany) on the day before the experiment, to avoid induced synchronicity of the cell cycle after splitting. In a typical experiment to follow cell cycle dynamics, cells were monitored from 48 to 120 hrs, with images taken every 10 min. In experiments where positive feedback at mitotic entry was perturbed using the WEE1 inhibitor PD166285 (Sigma-Aldrich, MO, USA) at 3 µM/mL, and DMSO as control, cells were imaged for 5 hrs after adding the inhibitor (or DMSO), with images taken every 4 min with the same acquisition conditions.

Live-cell imaging was performed on either an inverted motorized Leica Timelapse DMI6000B (Leica Microsystems) or in an inverted motorized Nikon ECLIPSE TI widefield microscope (Nikon) equipped with the Hamamatsu Orca-FLASH4.0 sCMOS (C11440, Hamamatsu) or Andor iXon Ultra 888 EMCCD (Andor) camera, respectively, both with temperature, humidity, and CO_2_ controls to maintain the sample integrity. Leica Timelapse DMI6000B (Leica Microsystems) images were acquired with an HCX PL FLUOTAR L 20x/0.40 Corr Ph1, using a binning 2×2 and a 16-bit depth, exposure time of 20ms with a lamp intensity of 20. Images were acquired every 10 min for 72 hrs. The microscope was controlled through the LAS X software (version 3.8.1.26810) using the navigation tool “LAS X Navigator”.

When using the Nikon ECLIPSE TI, fluorescence images were acquired with a Plan Apo λ 20x/0.75 DIC N2 objective. To excite the YFP the Lumencor Spectra X Teal – 510/25 nm was used in combination with the following filter set cube: CFP/YFP - ex 416/501 - bs 440/520 - em 464/547, using a LED % of 10 and an exposure time of 150ms. For mCherry, the Lumencor Spectra X Green – 550/15 was used with LED % of 15, an exposure time of 200ms, in combination with the following filter set cube:DAPI/FITC/TRITC/Cy5 – Pinkel Quad (FF410/504/582/669/Di01). Brightfield images were acquired with exposure time of 10ms with a lamp intensity of 5V. Images were acquired using binning 2×2 and a 16-bit depth.

### Live-cell and fixed-cell correlative microscopy

HDFs were grown in μ-Slide 2 Well ibiTreat (Ibidi GmbH, Gra□felfing, Germany) and analysed accordingly to the experimental layout shown in Figure 6A. Images were acquired on a Leica Timelapse DMI6000B (Leica Microsystems) with an HCX PL FLUOTAR L 20x/0.40 Corr Ph1, using a binning 2×2 and a 16-bit depth, exposure time of 20ms with a lamp intensity of 20. Neighbour fields (40-46) were imaged every 4 min for 48hrs. Following the long-term live-cell imaging, cells were processed for 53BP1/p21 and p16 double immunostaining as described in previous sections and the same neighbour fields were acquired using a ×40 LD/0.6 NA objective. LAS X software (Leica Microsystems) was used for image acquisition and ImageJ/Fiji software for image analysis.

### Real-time quantitative PCR

Total RNA was extracted using TRIzol reagent (Invitrogen, CA, USA), treated with DNAse I (Thermo Scientific), and precipitated with sodium acetate in ethanol. iScript Synthesis Kit (Bio-Rad, CA, USA) was used for cDNA synthesis. RT-qPCR was performed with iTaq Universal SYBR Green Supermix (Bio-Rad, CA, USA) in a CFX96 Touch Real[Time PCR Detection System (Bio-Rad, CA, USA) according to manufacturer’s instructions. Data were analysed using the CFX Maestro software (Bio[Rad, CA, USA). Three technical replicates were used per target gene. Expression was normalized to the *TBP* and *HPRT1* housekeeping genes, and different experimental samples were normalized to the mean expression of the control samples.

### Western blotting

HDFs were detached with Trypsin, washed twice with ice-cold PBS, and lysed in lysis buffer (150 nM NaCl, 10 nM Tris-HCl pH 7.4, 1 nM EDTA, 1 nM EGTA, 0.5% IGEPAL) with 1x cOmplete, EDTA-free Protease Inhibitor (Roche, Mannheim, Germany). Lysates were quantified for protein content by the Lowry Method (DCTM Protein Assay, Bio-Rad, CA, USA). Twenty micrograms of total extract were loaded in SDS-polyacrylamide gels for electrophoresis and transferred onto nitrocellulose membranes for western blot analysis. Membranes were blocked during 1h with TBS (50 mM Tris-HCl, 150 mM NaCl) containing 5% low-fat milk. Primary antibodies were diluted in TBS containing 2% low-fat milk: rabbit anti-FOXM1 (#13147, ProteinTech Group, Inc, IL, USA), 1:1000; mouse anti-FZR1 (#SC-56312, Santa Cruz Biotechnology, CA, USA) 1:1000; and mouse anti-GAPDH (#T5168, Sigma-Aldrich, CA, USA), 1:50000. Horseradish peroxidase-conjugated secondary antibodies goat anti-rabbit (1:10000, 111-035-003, Jackson ImmunoResearch Europe Ltd., Ely, UK) and goat anti-mouse (1:15000, 115-035-003, Jackson ImmunoResearch Europe Ltd., Ely, UK) were diluted in TBS + 2% low-fat milk. Detection was done using Clarity Western ECL Substrate reagent (Bio-Rad, CA, USA) as per manufacturer’s instructions. Quantitative analysis of protein levels was carried out using a GS-900 calibrated densitometer with ImageLab software (Bio-Rad, CA, USA).

### Image analysis

Live-cell movies and fixed-cell experiments were blindly quantified using ImageJ/Fiji software ^58^.

### Statistical analysis

All experiments were repeated at least two times. Sample sizes and statistical tests used for each experiment are indicated in the respective figure captions. Data are shown as mean ± SD. GraphPad Prism version 8 was used to analyse all the data. Data were tested for parametric versus non-parametric distribution using D’Agostino–Pearson omnibus normality test. Mann-Whitney U test, two-tailed Fisher’s exact test and two-sided Kruskal-Wallis with Dunn’s multiple comparison correction tests were then applied accordingly to determine the statistical differences between different groups (\**p*<0.05, \*\**p*<0.01, \*\*\**p*<0.001, \*\*\*\**p*<0.0001, and *ns* not significant *p*>0.05).

## SUPPLEMENTARY FIGURE LEGENDS

**Supplementary Figure 1. Changes in cell cycle dynamics along aging.**

***A.*** Relative duration of cell cycle, G1-, S-, G2-phases and mitosis of neonatal, 10-, 21-, 42- and 87-year-old HDFs normalized to neonatal condition. All values represent Mean ± SD. n>50 cells, ≥2 independent experiments.

***B.*** Pseudo-timeline of the behaviour of cell cycle, G1-, S-, G2-phases and mitosis duration along aging using neonatal, 10-, 21-, 42- and 87-year-old HDFs. All values represent mean ± SD. The trend lines, R-squared (*R^2^*) and p-value are shown. n>50 cells, ≥2 independent experiments.

***C.*** Duration of mitosis as a function of G1-phase, duration of S-phase as a function of G1-phase and duration of G2-phase as a function of S-phase in neonatal (dark blue), 10- (light blue), 21- (yellow), 42- (orange) and 87-year-old (red) HDFs expressing PIP-FUCCI. The trend lines are shown. n>60 cells, ≥2 independent experiments.

***D.*** Duration of mitosis measured by the time between nuclear envelope breakdown and nuclear envelope reformation as a function of cell cycle length in neonatal, 10-, 21-, 42- and 87-year-old HDFs by phase contrast. The trend lines, Pearson coefficient (*r*) and p-value are shown. n>60 cells, ≥2 independent experiments.

**Supplementary Figure 2. Older cells are less prone to commit to a new division cycle.**

***A.*** Schematic of the experimental design to test whether positive feedback at mitotic entry is deregulated during aging. The hypothesis is that the sharp sigmoidal activation of CDK1 at mitotic entry (left) is lost as the cell ages (right).

***B.*** Bar plot of neonatal cells representing the time since last division (grey bars) until treated with DMSO (left, black) or with Wee1 inhibitor (right, red). Filled bars represent no entering into mitosis within 5 hours of treatment with either DMSO or Wee1 inhibitor. Open bars represent cells that entered mitosis (end of the bars represents the time of mitotic entry). Box plot representing the time since last division of neonatal cells treated with DMSO (black) and Wee1 inhibitor (red) that did not divide and that divided (≥2 independent experiments, >50 cells).

***C.*** Bar plot of 87-year-old cells representing the time since last division (grey bars) until treated with DMSO (left, black) or with Wee1 inhibitor (right, red). Filled bars represent no entering into mitosis within 5 hours of treatment with either DMSO (black) or Wee1 inhibitor (red). Open bars represent cells that entered mitosis. Box plot representing the time since last division of 87-year-old cells treated with DMSO (black) and Wee1 inhibitor (red) that did not divide and that divided (≥2 independent experiments, >50 cells).

**Supplementary Figure 3. Modulation of FOXM1 levels impacts on mitosis and cell cycle length.**

***A.*** Gene expression of mitotic entry feedback loop players between neonatal and 87- year-old HDFs mitotic extracts, normalized to neonatal samples ^19^.

***B.*** Western blot analysis and quantification of FOXM1 protein levels following RNAi depletion in neonatal HDFs. GAPDH was used as loading control. All values represent mean ± SD. 4 independent experiments. * *p*≤0.05 by Mann-Whitney U test.

***C.*** Western blot analysis and quantification of FOXM1 endogenous protein levels in 87-year-old cells transduced with lentiviruses to overexpress a truncated form of FOXM1 (FOXM1-dNdK) ^19^ in the absence (-Dox) or the presence of doxycycline (+Dox). GAPDH was used as loading control. All values represent mean ± SD. 6 independent experiments. ** *p*≤0.01 by Mann-Whitney U test.

***D.*** Mitotic duration (time between NEB and NER) and cell cycle length measured in phase-contrast movies by the time between two consecutive mitoses in neonatal HDFs, siFOXM1-depleted neonatal HDFs, 87-year-old HDFs and 87-year-old HDFs overexpressing FOXM1. All values represent mean ± SD. n>50 cells, ≥2 independent experiments. *ns* nonsignificant, ** *p*≤0.01, *** *p*≤0.001, **** *p*≤0.0001 by two-sided Kruskal-Wallis with Dunn’s multiple comparison correction.

**Supplementary Figure 4. Cdh1 levels increase with advancing aging and correlate with decreased cell proliferative fitness.**

***A.*** Representative images of Cdh1 intensity levels by immunofluorescence in neonatal, 10-, 21-, 42- and 87-year-old HDFs. Scale bar, 20μm.

***B.*** Representative images of Cdh1 intensity levels by immunofluorescence mock- and siFOXM1-depleted neonatal cells. Scale bar, 20μm.

***C.*** Quantification of Cdh1 intensity levels by immunofluorescence analysis in control (OECdh1 -Dox) or overexpressing Cdh1 (OECdh1 +Dox) neonatal HDFs. Levels were normalized to control. All values represent mean ± SD. n≥550 cells, 6 independent experiments. *** *p*≤0.001 by Mann-Whitney U test.

***D.*** Quantification of Cdh1 positive cells by immunofluorescence analysis in control or overexpressing Cdh1 neonatal HDFs. Levels were normalized to control. All values represent mean ± SD. n≥1100 cells, 5 independent experiments. ** *p*≤0.01 by two-tailed Fisher’s exact test.

***A. E.*** Quantification of FOXM1 intensity levels by immunofluorescence analysis in control or overexpressing Cdh1 neonatal HDFs. Levels were normalized to control. All values represent mean ± SD. n≥1100 cells, 6 independent experiments. *** *p*≤0.001 by Mann-Whitney U test.

***B. E.*** Percentage of mitotic cells (positive for phospho-histone 3 immunostaining – pH3+) in control or overexpressing Cdh1 neonatal cultures. All values represent Mean ± SD. n3≥000 cells, ≥2 independent experiments. **** *p*≤0.0001 by two-tailed Fisher’s exact test.

***C. F***. Quantification of KI67+ proliferating cells in control or overexpressing Cdh1 neonatal cultures. All values represent Mean ± SD. n≥3000 cells, 3 independent experiments. **** *p*≤0.0001 by two-tailed Fisher’s exact test.

**Supplementary Figure 5. siRNA-mediated depletion of *FZR1*/Cdh1 increases cell proliferation and cell cycle fitness in older cells.**

***A.*** Western blot analysis and quantification of Cdh1 protein levels following RNAi depletion in 87-year-old HDFs. GAPDH was used as loading control. All values represent mean ± SD. 4 independent experiments.

***B.*** Gene expression analysis of *FZR1*/Cdh1 by RT-qPCR in mock- or siCdh1-depleted 87y-year-old HDFs. All values represent mean ± SD.

***C.*** Representative images and quantification of FOXM1 intensity levels by immunofluorescence analysis in mock or siCdh1-depleted 87y-year-old HDFs. Levels were normalized to mock condition. All values represent mean ± SD. n>800 cells, 6 independent experiments. **** *p*≤0.0001, by Mann-Whitney U. Scale bar, 20μm.

***D.*** Cumulative percentage of dividing cells as a function of time following treatment with mitotic trigger in siCdh1-depleted 87y-year-old HDFs. Cells were either treated with DMSO (black) or with Wee1 inhibitor (Wee1 inh) (red) (n>250 cells, >3 independent experiments).

***E.*** Percentage of mitotic cells (pH3+) in mock- or siCdh1-depleted 87y-year-old HDFs. All values represent mean ± SD. n≥4000 cells, 4 independent experiments. **** *p*≤0.0001 by two-tailed Fisher’s exact test.

***F.*** Quantification of ki67+ proliferating cells in mock- or siCdh1-depleted 87y-year-old HDFs. All values represent mean ± SD. n≥3000 cells, 3 independent experiments.

**** *p*≤0.0001 by two-tailed Fisher’s exact test.

**Supplementary Figure 6. Mitotic stopwatch complex players are downregulated in aged mitotic HDFs.**

***A.*** Gene expression of mitotic stopwatch complex players in neonatal and 87-year-old HDFs mitotic extracts, normalized to neonatal ^19^.

***B.*** Correlation between relative p21 intensity levels of daughter cells and their time spent in interphase. The trend lines, Pearson coefficient (*r*) and p-values are shown. n>60 cells.

***C.*** Correlation between relative p16 intensity levels of daughter cells and their time spent in interphase. The trend lines, Pearson coefficient (*r*) and p-values are shown. n>60 cells.

## ACKNOWLEDGMENTS

We acknowledge Silvia Santos and Jorge Ferreira for kindly providing the vectors pCSII-EF1a-CyclinB1-venus and pLenti-PGK-Neo-PIP-FUCCI, respectively; and Nádia Santos for assisting with logistic equation analyses. We thank the personnel at i3S Scientific Platforms for technical support: CCGEN (P. Magalhães and T. Meireles), advanced light microscopy (P. Sampaio and M. Azevedo) and Translational Cytometry (C. Meireles and E. Cardoso). ALM is member of the Portuguese Platform of Bioimaging (PPBI-POCI-01-0145-FEDER-022122). We are grateful to J.C. Macedo and all laboratory members for their support.

## AUTHOR CONTRIBUTIONS

ARA and EL contributed to the conceptualization, development and supervision of the work. ARA and FG-S performed all wet lab experiments and respective analyses. ARA and EL were involved in funding acquisition. ARA and EL wrote the manuscript with input from FG-S. All authors have read and agreed to the final version of the manuscript.

