## Supplemental figures for "Feedback regulation between FOXM1 and APC/C^Cdh1^ determines the changes in cell cycle dynamics during aging"

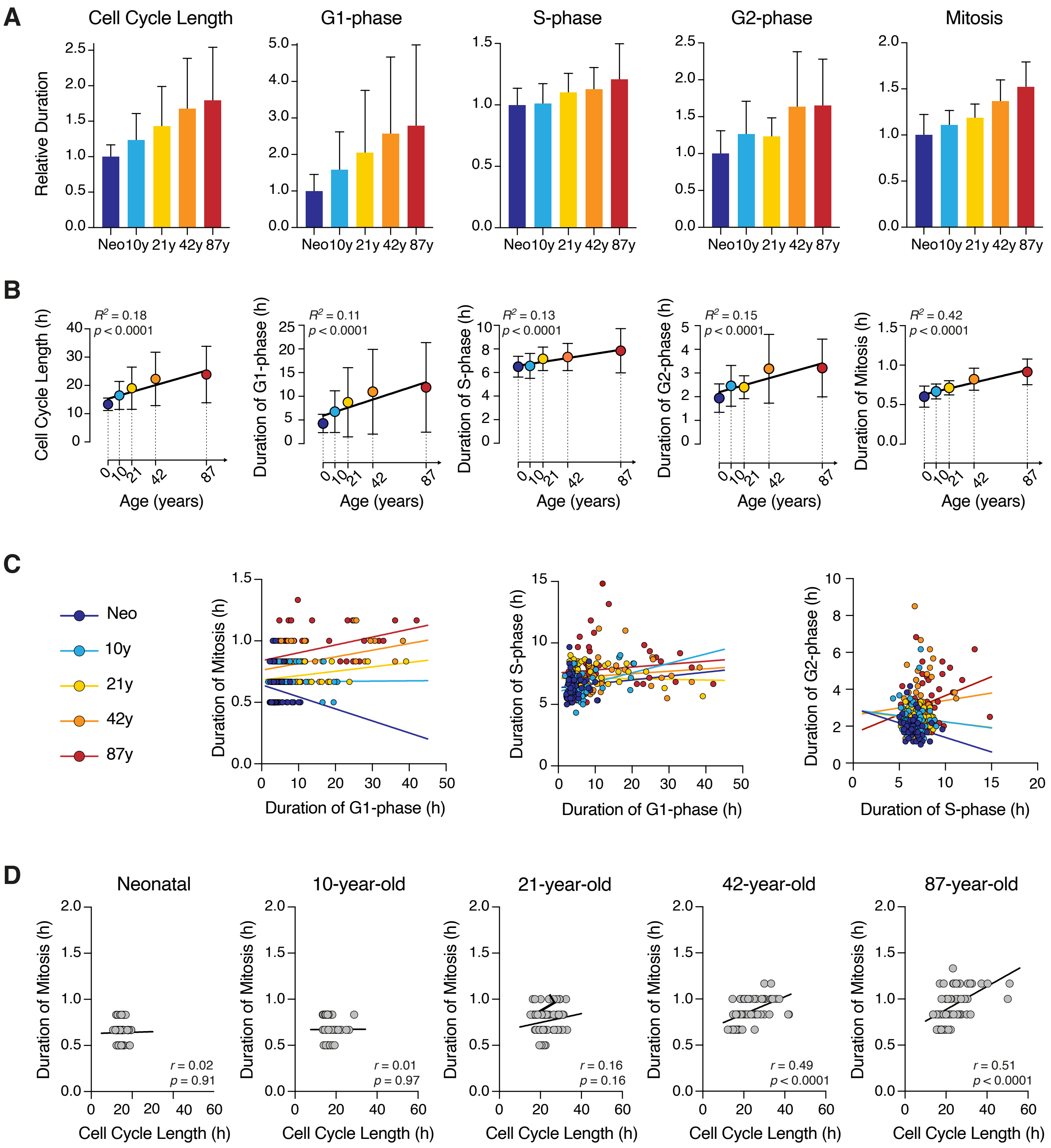


Araujo et. al 2025
Supp Figure 1

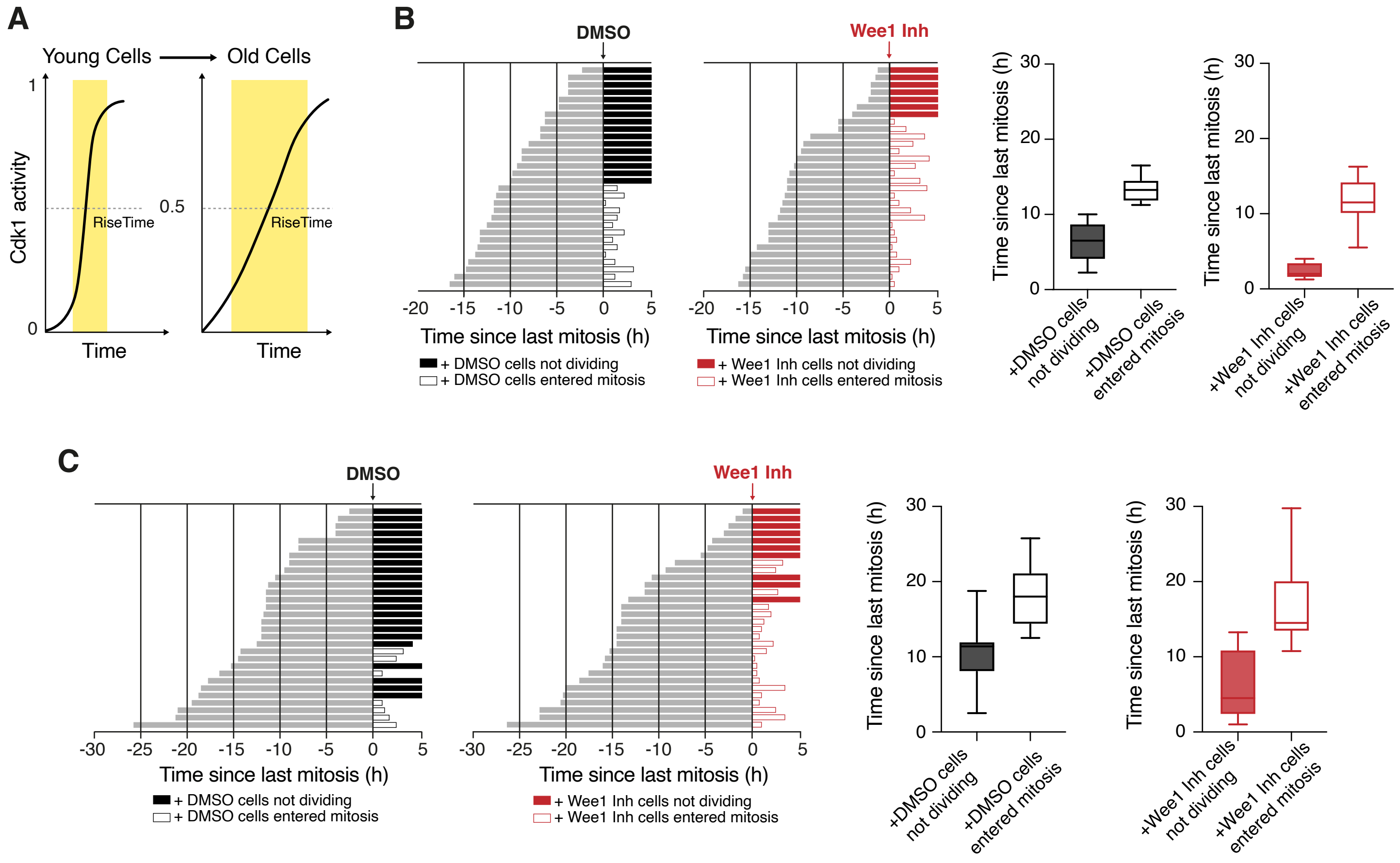


Araujo et. al 2025
Supp Figure 2

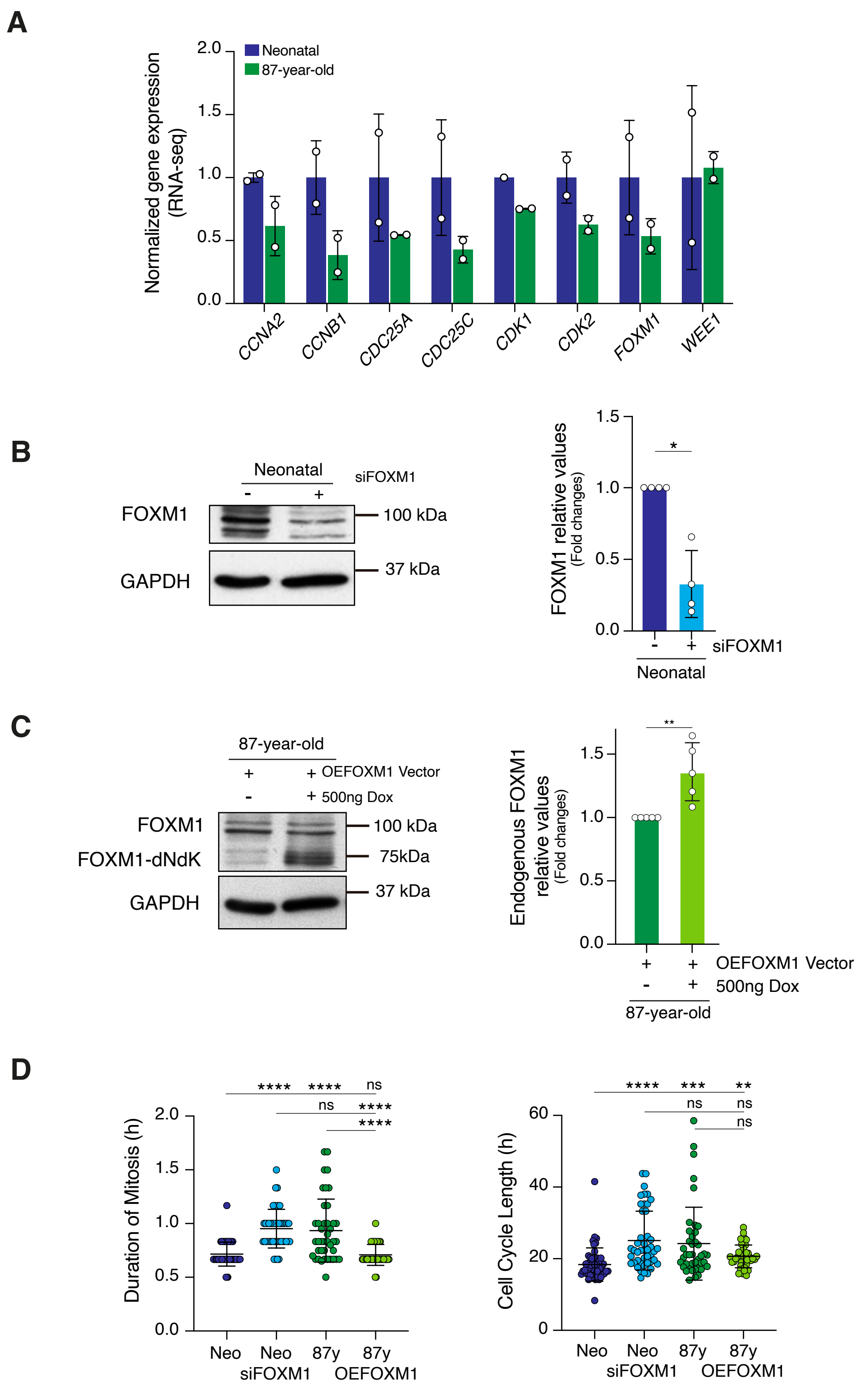


Araujo et. al 2025
Supp Figure 3

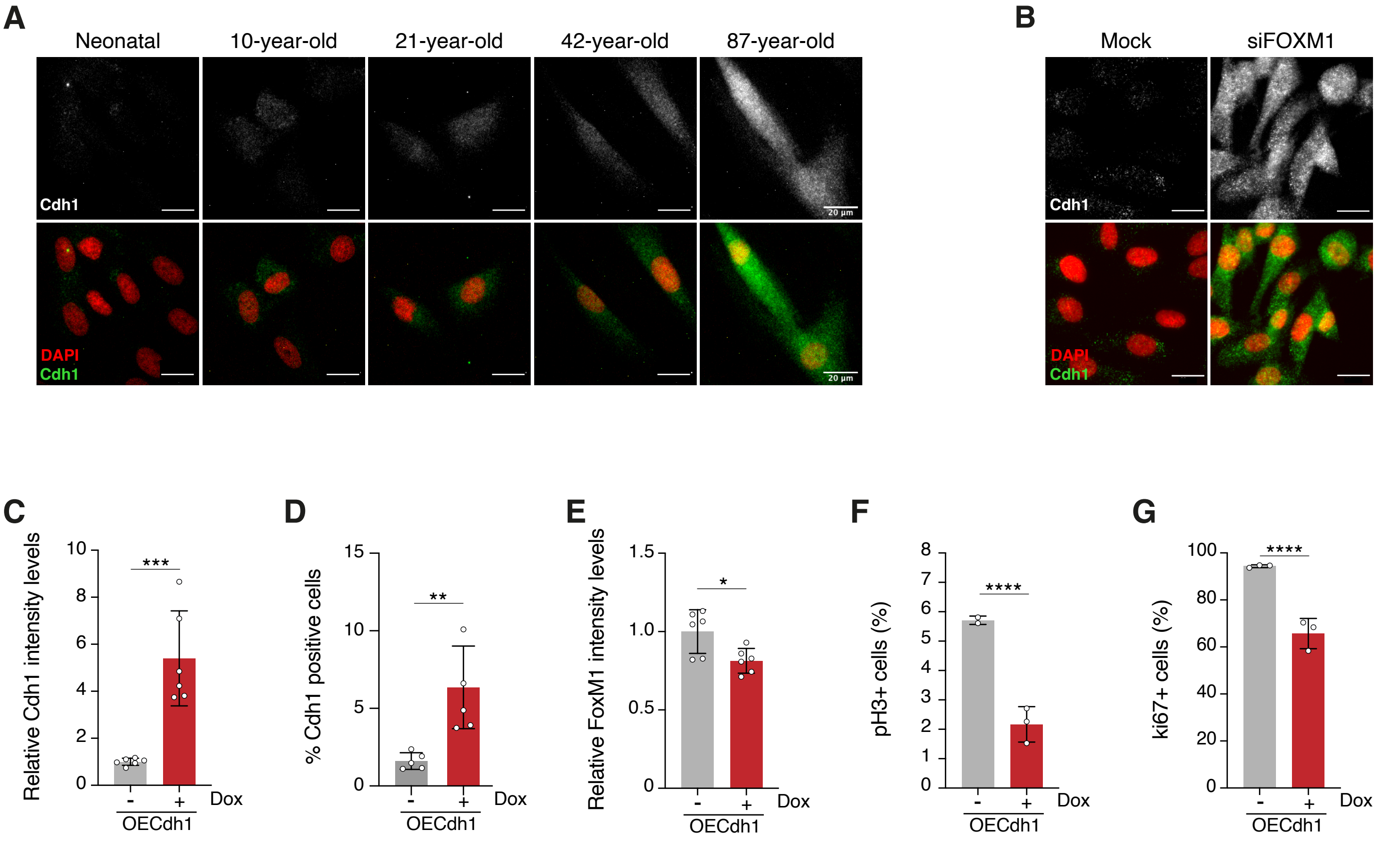


Araujo et. al 2025
Supp Figure 4

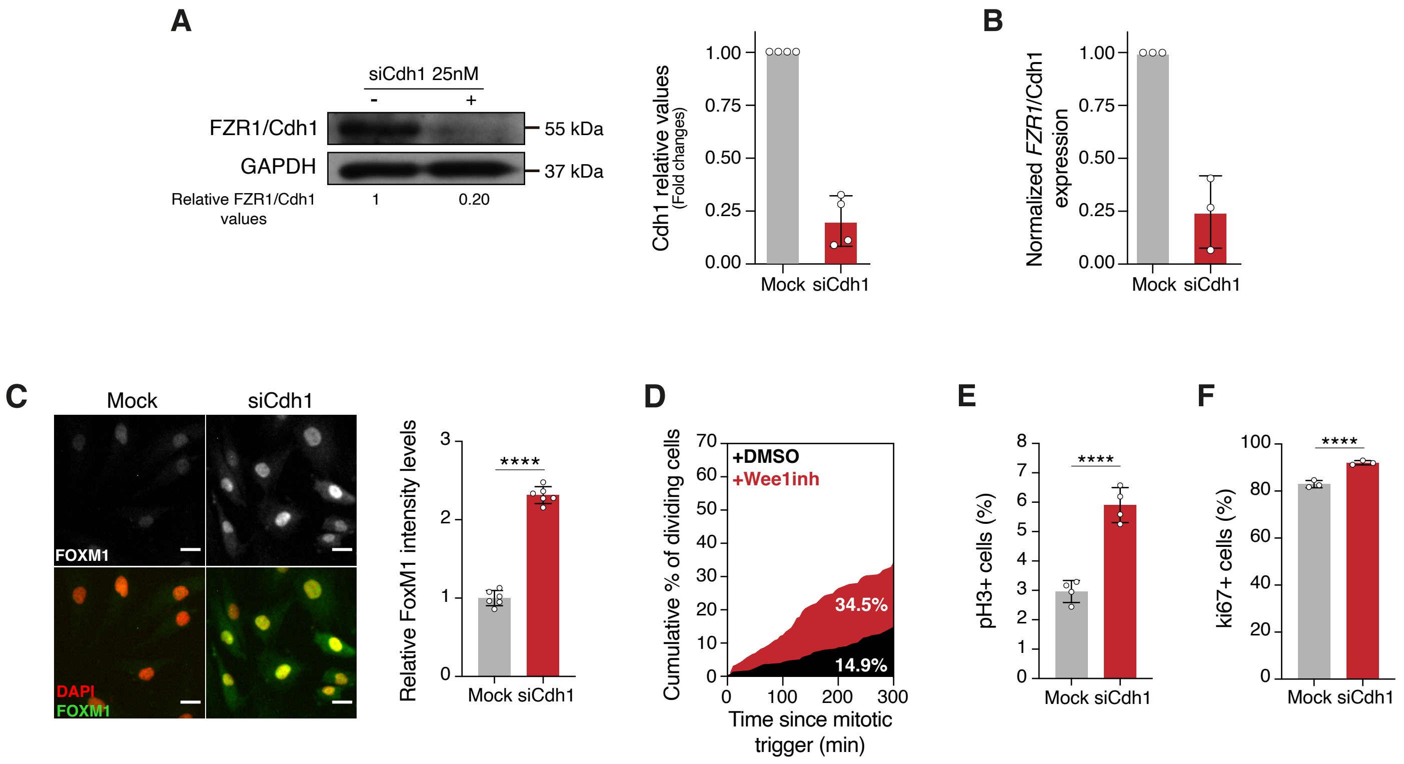


Araujo et. al 2025
Supp Figure 5

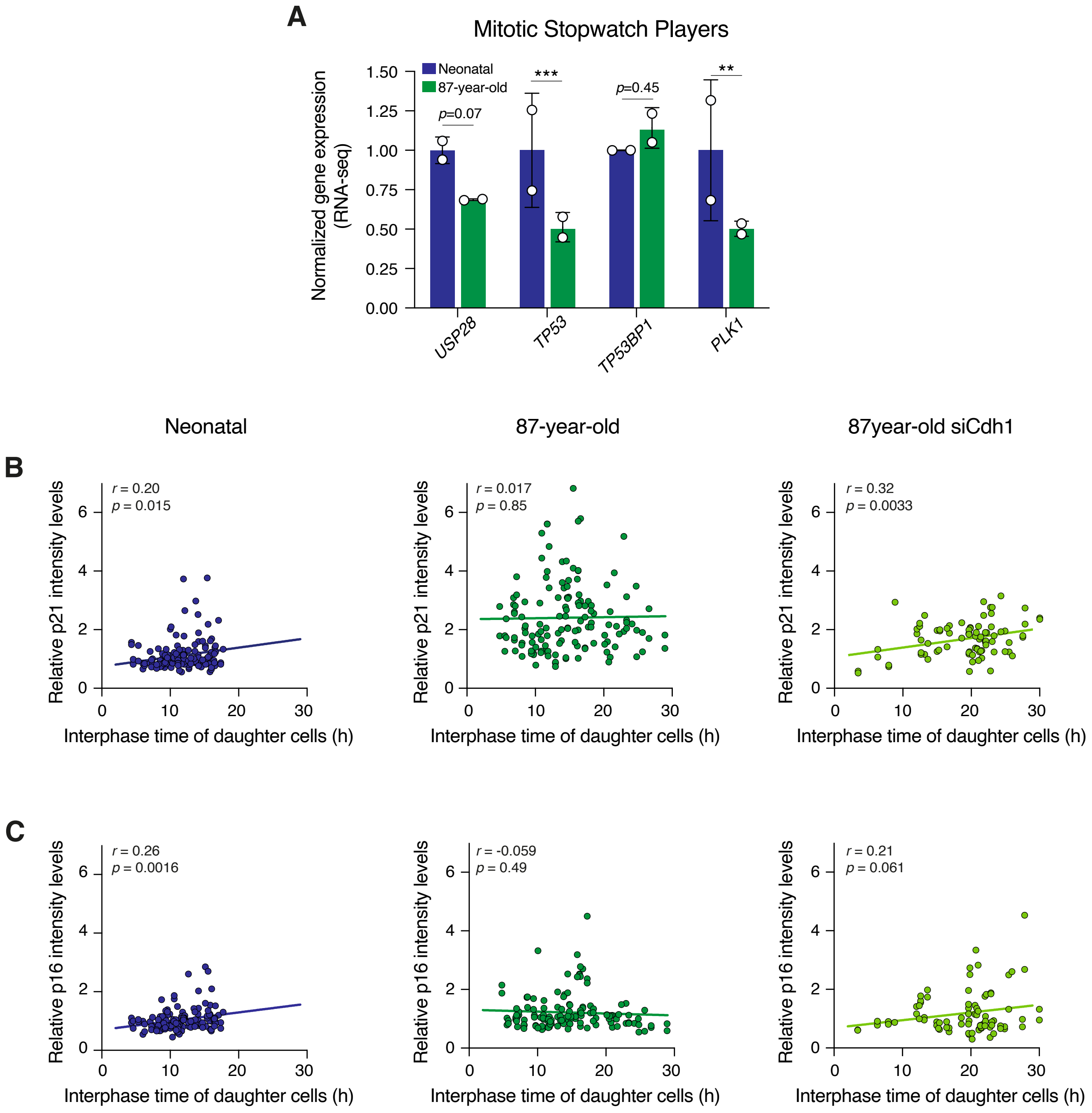


Araujo et. al 2025
Supp Figure 6
